# The DNA binding domain of the *Vibrio vulnificus* SmcR transcription factor is flexible and recognizes diverse DNA sequences

**DOI:** 10.1101/2020.10.30.362368

**Authors:** Jane D. Newman, Meghan M. Russell, Giovanni Gonzalez-Gutierrez, Julia C. van Kessel

## Abstract

The quorum-sensing regulon in vibrios is controlled by the LuxR/HapR family of transcriptional regulators. In *Vibrio vulnificus*, this regulator is called SmcR, and it controls expression of numerous virulence behaviors, including biofilm formation and elastase production. The consensus binding site of *Vibrio* LuxR/HapR/SmcR proteins is palindromic, as is common for regulators that bind as dimers with helix-turn-helix N-terminal DNA binding domains. However, the LuxR/HapR/SmcR consensus site is highly degenerate and asymmetric with variations in sequence at each promoter. To determine the mechanism of DNA site recognition, we generated separation-of-function mutants of SmcR that either repress or activate transcription but not both. The SmcR N55I protein is defective at transcription activation due to loss of binding to most DNA binding sites in activated promoters but retains interaction with RNA polymerase (RNAP) alpha. SmcR S76A, L139R, and N142D are defective for interaction with RNAP alpha but retain functional DNA binding activity. Using X-ray crystallography, we show that the wild-type SmcR dimer and the three RNAP-interaction mutants exhibit two conformations of the helix-turn-helix DNA binding domain. Conversely, the SmcR N55I X-ray crystal structure is limited to only one conformation and is restricted in recognition of single base-pair variations in DNA binding site sequences. These data support a model in which two mechanisms drive SmcR transcriptional activation: interaction with RNA polymerase and a multi-conformational DNA binding domain that permits recognition of variable DNA sites. Thus, the LuxR/HapR/SmcR proteins balance specificity for quorum-sensing targets and diversity to accommodate binding at hundreds of sites genome-wide.

**Significance:** The cell-cell communication system called quorum sensing controls expression of genes required for virulence in *Vibrio* bacteria species, including the potent human pathogen *Vibrio vulnificus*. The master transcriptional regulator of quorum-sensing genes in vibrios belongs to the LuxR/HapR/SmcR family. These regulators directly activate and repress transcription of >100 genes via binding to degenerate sites in promoter regions. We used X-ray crystallography to determine the structure of mutant SmcR proteins. Our experiments reveal that SmcR recognizes diverse sequences via a DNA binding domain that samples multiple conformations to accommodate variations in palindromic DNA sequences. Significantly, the DNA binding domain of SmcR is completely conserved in LuxR/HapR/SmcR family proteins, suggesting that this mechanism is representative of quorum-sensing regulation in other vibrios.

## Introduction

Bacterial pathogens oscillate between vastly different environments outside and within their host organism. Cells must tune gene expression to respond to changes such as nutrient acquisition, temperature, pH, salt, and the presence of other microorganisms. Upon entering a host, bacteria need to upregulate genes required for colonization and establishing an infection, including cell surface appendages for attachment to surfaces, secretion systems for delivery of toxins, and enzymes for digestion of macromolecules for nutrient acquisition (1–5). Vibrios are Gram-negative bacteria that are found in the marine environment, and many of these bacteria are also pathogens of a variety of fish and shellfish (6–10). Notably, some vibrios are potent human pathogens such as *Vibrio cholerae, Vibrio parahaemolyticus*, and *Vibrio vulnificus*. In vibrios, one major regulatory system that controls expression of virulence genes is quorum sensing.

Quorum sensing is a process of cell-cell signaling that allows bacteria to control group behaviors in response to an increase in population density (11). Quorum-sensing transcription factors are responsible for regulating hundreds of genes. In *Vibrio* species, the master quorum-sensing transcriptional regulators are the LuxR/HapR/SmcR-type proteins that are produced at high cell density (12, 13). In vibrios, these proteins are the central regulators of virulence gene expression, including LuxR in *Vibrio harveyi*, HapR in *V. cholerae*, OpaR in *V. parahaemolyticus*, and SmcR in *V. vulnificus*. Indeed, deletion or inhibition of *smcR* in *V. vulnificus* decreases pathogenesis in both mouse and shrimp models (14, 15). Thus, these core regulatory proteins are central to our understanding of the influence of quorum sensing on virulence.

Transcriptional activation in bacteria canonically occurs via interactions with RNA polymerase (RNAP) via recruitment or stabilization of RNAP interactions with DNA. Conversely, repression generally occurs through blocking key promoter elements or binding of required activator proteins (16, 17). The complexity of these processes in bacteria has recently become more fully appreciated with the advent of deep-sequencing technologies and the ability to connect transcriptomics to nucleoprotein complexes at promoters (18). Such experiments have revealed numerous important aspects of the mechanism of regulation by the LuxR/HapR/SmcR family of transcription factors. These proteins are completely different from the LuxI/LuxR quorum-sensing systems that synthesize and directly bind autoinducers, respectively, to enable DNA binding and regulation. Rather, LuxR/HapR/SmcR are sub-members of the broader TetR family of transcription factors. Previous work has shown that these quorum-sensing regulators are distinct from most TetR proteins because they are able to activate and repress transcription of many genes, while canonical TetR proteins only repress a few genes (19). Similar to most TetR proteins, the binding consensus sequences for LuxR/HapR/SmcR proteins tends to be between 20 to 22 base pairs long (20–22). In contrast to typical TetR proteins, the site for LuxR/HapR/SmcR is degenerate and asymmetric (20, 22–24). The diverse binding sites of the LuxR/HapR/SmcR group also have a wide array of binding affinities, with dissociation constants ranging from approximately 0.5-100 nM (23). Though a significant portion of biochemistry has been performed with the founding member of this family, LuxR, the SmcR protein is highly amenable to structural studies by X-ray crystallography. The HapR protein has also been amenable to crystallography (25), but the DNA-binding domain of LuxR and SmcR are 100% identical whereas there are some differences in HapR (Fig. S1A), making the study of SmcR more appealing for studying DNA-binding mechanisms. Thus, we focus on the mechanisms of transcriptional activation by the *V. vulnificus* TetR-type protein SmcR.

Among vibrios, *Vibrio vulnificus* is considered a potent human pathogen due to a mortality rate of greater than 50% in primary septicemia infection cases (26). In *V. vulnificus*, the master quorum-sensing transcription factor SmcR is responsible for activating virulence genes via many factors, one of which is by increasing the expression of an elastase gene *vvpE* by directly binding to the promoter (27) It is hypothesized that this activation mechanism functions via SmcR interaction with RNAP. A direct interaction between LuxR in *Vibrio harveyi* has been shown *in vitro* and *in vivo* (28), and this interaction with RNAP likely also exists with SmcR (29). We must note that *Vibrio harveyi* BB120 has been reclassified as *Vibrio campbellii* BB120 (a.k.a. ATCC BAA-1116), but we refer to this organism in this manuscript as *V. harveyi* for consistency with the literature.

In this study we use structural and biochemical approaches to investigate SmcR-DNA and SmcR-protein interactions and the impact of these functions on activation of quorum-sensing genes. Our data show that a specific amino acid substitution in the DNA binding domain of SmcR renders the protein incapable of binding DNA at most established binding sites, likely by limiting the conformational states of the helix-turn-helix that interacts with DNA. Conversely, amino acid substitutions in the RNAP interaction domain of SmcR do not alter the conformations of the DNA binding domains. Our results support a model that multiple conformations of SmcR allow recognition of diverse DNA binding sequences in activated quorum-sensing promoters.

## Results

### The N55I substitution in SmcR separates activation and repression phenotypes

The LuxR/HapR/SmcR proteins are of interest because they both activate and repress transcription, which is unique among the TetR family of transcription factors. Previously, genetic screens have been performed to find separation-of-function mutants of LuxR in order to study the mechanisms of activation or repression individually (23). Substitutions of specific amino acids in the DNA binding domain of LuxR in *V. harveyi* (*e.g.,* N55I; Fig. 1A) render the protein unable to bind to specific DNA binding sites, thus resulting in lack of either activation or repression of *V. harveyi* promoters (23). In addition, substitutions in alpha helices 4 and 7 of LuxR (S76A, N142D, L139R) result in decreased or eliminated activation activity due to loss of interaction(s) with RNAP (Fig. 1A) (28). Because SmcR shares 100% amino acid identity with LuxR in the DNA binding domain, shares 92% identity overall, and contains the same critical residues in the RNAP interaction domain (Fig. S1A), we hypothesized that substitutions in the same positions in SmcR would yield similar activation- and repression-specific phenotypes. We specifically focused on SmcR N55I for several reasons: 1) the S76A, N142D, and L139R substitutions were shown to have the same *in vivo* phenotypes in both LuxR (*V. harveyi*) and HapR (*V. cholerae*), indicating that the interaction with RNAP alpha is conserved across vibrios (28), 2) the LuxR N55I protein is defective at DNA binding activity at specific sequences but not all known binding sites (23), 3) the LuxR N55I substitution had the largest decrease in activation activity while retaining wild-type repression activity (23), 4) the asparagine is in a region of the DNA binding domain that is completely conserved in all LuxR/HapR-type proteins but not *Escherichia coli* TetR, and 5) substitution of N55 with four other amino acids (Y, K, S, A) all resulted in loss of activation (19, 23). We assayed function of SmcR N55I using a previously published dual-promoter reporter plasmid containing the activated promoter P_*luxC*_ driving expression of *gfp* and the repressed promoter P_*05222*_ driving expression of *mCherry* (23). As we predicted, the N55I substitution in SmcR resulted in complete loss of activation of the *luxC* promoter, whereas repression was maintained and even increased compared to wild-type SmcR (Fig. 1B).

**Figure 1.**
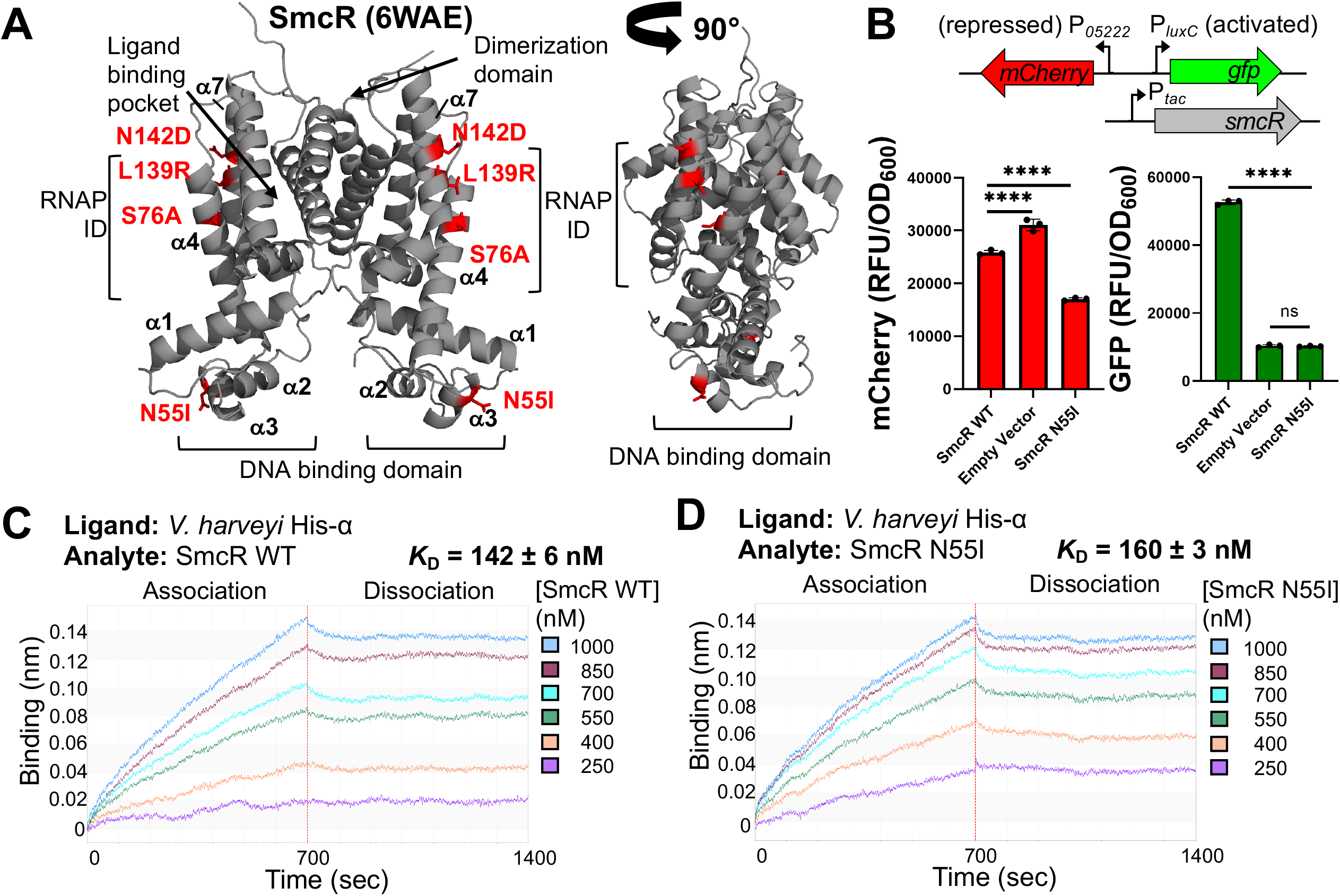
The SmcR N55I substitution mutant is deficient in transcriptional activation but not RNAP interaction. **(A)** X-ray structure of SmcR (PDB: 6WAE) in gray, with relevant substitution mutant residues (N55I, S76A, L139R, N142D) highlighted in red sticks. RNAP ID, RNA polymerase interaction domain. Alpha helices are indicated with numbers. **(B)** Fluorescence (GFP/OD_600_ or mCherry/OD_600_) was measured from biological triplicates of *E. coli* strains containing two plasmids: 1) a dual reporter plasmid pJV064 containing P_*05222*_-mCherry (repressed promoter) and P_*luxC*_-GFP (activated promoter), and 2) a plasmid expressing IPTG-inducible copies of either *smcR* (pJN22), the substitution mutant *smcR N55I* (pJN27), or an empty vector control (pMMB67EH-kanR). Asterisks (****) indicate that the fluorescence values are significantly different than the wild-type counterpart (*p* < 0.0001; two-way analysis of variance (ANOVA), followed by Tukey’s multiple-comparison test; *n* = 3). **(C, D)** BLI analysis of binding reactions containing 200 nM *V. harveyi* His-RNAP α (ligand) and the indicated concentrations of analyte wild-type SmcR (C) or SmcR N55I (D). The average calculated binding affinities with standard deviations (*K*_D_; *n* = 3) are listed.

Our previous work with LuxR/HapR had revealed the RNAP interaction domain on LuxR and established that LuxR interacts with RNAP alpha (28). We used biolayer interferometry (BLI) to show that SmcR also interacts with RNAP alpha (*K*_D_ = 142 +/− 6 nM, Fig. 1C) with a similar affinity to LuxR (*K*_D_ = 139 +/− 19 nM). This is the first definitive *in vitro* experiment showing that SmcR and RNAP directly interact. We also observed that SmcR N55I interacts with RNAP alpha (*K*_D_ = 160 +/− 3 nM) with an affinity similar to wild-type SmcR (Fig. 1D). From these data, we conclude that the N55I substitution in the DNA binding domain of SmcR results in a loss of transcription activation that is not caused by a disruption in the RNAP alpha-interaction domain on SmcR.

### SmcR N55I has limited DNA binding site recognition

To examine the diversity of sequences bound by SmcR N55I, we assayed DNA binding activity at the eight binding sites A-H in the *luxC* promoter from *V. harveyi* using electrophoretic mobility shift assays (EMSAs). This promoter is ideal for this experiment because the LuxR binding sequence of each of these sites has been determined, and SmcR binds to each of these sites with similar affinity to LuxR (Fig. 2A, 2B, S2A) (30). SmcR N55I only binds well to sites B and H, at which it retains near wild-type levels of DNA binding (Fig. 2B, S2A, S2B, Table 1). The deficient binding of SmcR N55I to six out of eight P_*luxC*_ binding sites correlates to the lack of SmcR N55I activation activity at the *luxC* promoter *in vivo* (Fig. 1B). We note that some EMSA DNA substrates (*e.g.*, sites D and E) migrate differently at high concentrations of SmcR protein that produce quantifiable DNA shifts (Fig. S2A). These high-migrating substrates do not match the migration pattern of single-dimer shifted bands (*e.g.*, sites A, B, C, F, G, H; Fig. S2A), and these substrate-specific shifted bands are also observed with LuxR (30). Sites D and E overlap by six basepairs (Fig. S1B) (30). For each site, we used substrates that contained mutations in the other binding site that we have shown eliminate or greatly decrease binding of LuxR (Fig. S1B). However, we suspect that these high-migrating bands comprise the binding of a second SmcR dimer to the mutual half-site that is still intact, which is why they are only observed for sites D and E and only at high concentrations that we previously did not test. Another hypothesis is that upon substrate binding SmcR may alter the DNA conformation in a way that changes the migration pattern (*e.g.*, DNA bending).

**Table 1.** EMSA analyses of SmcR protein-DNA interactions.

| DNA Substrate | Oligos | SmcR Protein | $K_d$ (nM) |
| --- | --- | --- | --- |
| $P_{vvpE}$ | AB324/AB325 | WT | 0.1295 |
| $P_{vvpE}$ | AB324/AB325 | N55I | 303.5 |
| $P_{vvpE}$ | AB324/AB325 | S76A | 3.438 |
| $P_{vvpE}$ | AB324/AB325 | L139R | 8.163 |
| $P_{vvpE}$ | AB324/AB325 | N142D | 18.29 |
| $P_{vvpM}$ | AB316/AB317 | WT | 0.02788 |
| $P_{vvpM}$ | AB316/AB317 | N55I | 0.327 |
| $P_{vvpM}$ | AB316/AB317 | S76A | 0.2918 |
| $P_{vvpM}$ | AB316/AB317 | L139R | 1.023 |
| $P_{vvpM}$ | AB316/AB317 | N142D | 2.607 |
| $P_{luxC}$ Site B | JDN54/JDN55 | WT | 0.1311 |
| $P_{luxC}$ Site B | JDN54/JDN55 | N55I | 0.3222 |
| $P_{luxC}$ Site H | JCV369/JCV620* | WT | 0.009997 |
| $P_{luxC}$ Site H | JCV369/JCV620* | N55I | 0.07784 |
| $P_{luxC}$ Site E (abolish D binding) | RC212/RC213* | WT | 11.98 |
| $P_{luxC}$ Site E (abolish D binding) | RC212/RC213* | N55I | 240.6 |
| $P_{luxC}$ Site E<sup>T→A</sup> | JDN44/JDN45 | WT | 1.534 |
| $P_{luxC}$ Site E<sup>T→A</sup> | JDN44/JDN45 | N55I | 13.41 |
| $P_{luxC}$ Site E<sup>A→T</sup> | JDN46/JDN47 | WT | 2.714 |
| $P_{luxC}$ Site E<sup>A→T</sup> | JDN46/JDN47 | N55I | 44.61 |
| $P_{luxC}$ Site E<sup>T→A, A→T</sup> | JCV1107/JCV1108 | WT | 2.045 |
| $P_{luxC}$ Site E<sup>T→A, A→T</sup> | JCV1107/JCV1108 | N55I | 1.909 |
| $P_{luxC}$ Site A | JCV1075/JCV1076* | WT | 9.944 |
| $P_{luxC}$ Site C | JCV1079/JCV1080* | WT | 103.2 |
| $P_{luxC}$ Site D (abolish E binding) | RC224/RC225* | WT | 17.96 |
| $P_{luxC}$ Site F | JCV1087/JCV1088* | WT | 104.8 |
| $P_{luxC}$ Site G | JCV367/JCV619* | WT | 7.558 |

\*Indicates this oligonucleotide pair was published in a previous study (30).

**Figure 2.**
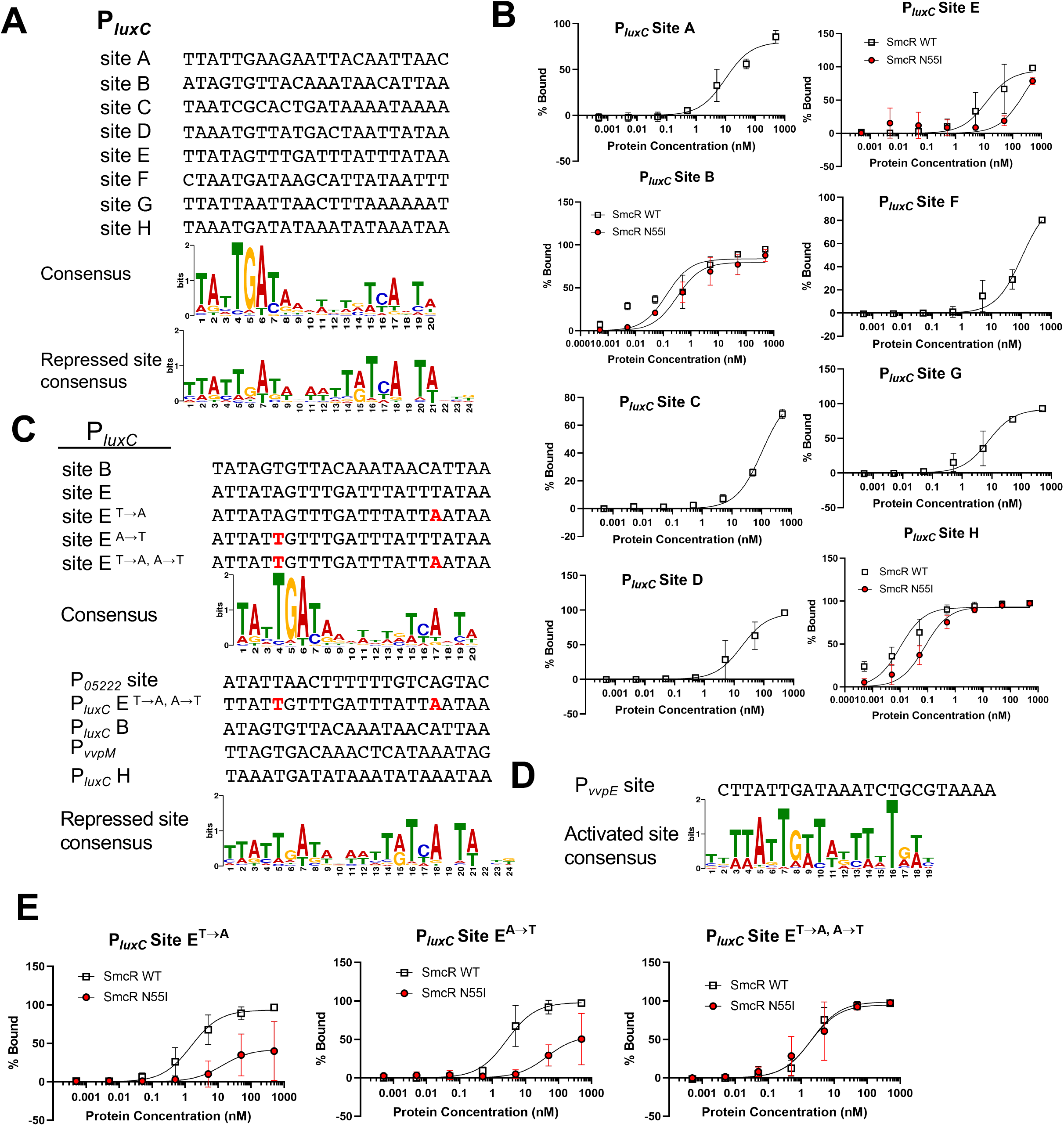
SmcR mutant N55I has limited DNA binding activity. **(A)** Alignment of P_*luxC*_ binding sites compared to the LuxR consensus binding sequence and the LuxR repressed binding site sequence, which was generated from the subset of LuxR repressed binding sites (23). **(B)** EMSA reactions consisting of 0.5 nM radiolabeled DNA substrates (P_*luxC*_ binding sites A through H, oligonucleotides listed in Table 1) and purified SmcR wild-type (white squares) or SmcR N55I protein (red circles) with increasing concentrations (0.0005, 0.005, 0.05, 0.5, 5, 50, 500 nM protein). Note that substrates for which no quantifiable shifts were observed for SmcR N55I do not have data points on graphs (Fig. S2A). Data points show the average of three independent experiments, and error bars indicate the standard deviation. **(C)** (top) Alignment of P_*luxC*_ binding sites B, E, and substitutions made in Site E compared to LuxR consensus motif and (bottom) alignment of P_*luxC*_ binding sites B, E^T→A, A→T^, H, P_*05222*_, and P_*vvpM*_ compared to the LuxR repressed consensus motif (23). Mutations in sites are indicated in red text. **(D)** P_*vvpE*_ binding sites aligned with the LuxR consensus motif for activated sites (23). **(E)** EMSAs comparing SmcR WT and SmcR N55I binding at P_*luxC*_ binding sites E^T→A^, E^A→T^, and E^T→A, A→T^ (oligonucleotides listed in Table 1 with increasing protein concentrations (0.0005, 0.005, 0.05, 0.5, 5, 50, 500 nM protein concentrations).

To examine differences between the sites, we focused on sites B and H because we observed that the SmcR N55I mutant has similar binding activity to wild-type at these sites (Fig. 2B, Table 1). Using the LuxR DNA binding site consensus as a guide, we aligned the sequences of LuxR binding sites in the *luxC* promoter to identify any sequence patterns that might contribute to binding differences between B and H and the other sites (Fig. 2C). We noted that several sites contained substitutions at highly conserved positions in the consensus, such as the T4 and A17 sites. To test the importance of these nucleotides, we introduced substitutions in the sequence of site E (A17→T or T4→A, or both) so that it more closely resembles site B and the consensus (Fig. 2C, 2E). The single nucleotide substitutions each increase wild-type SmcR binding to the substrates compared to wild-type site E but have little to no effect on SmcR N55I binding (Fig. 2E). Rather, both substitutions are required to restore SmcR N55I binding to wild-type levels at site E (Fig. 2E, Table 1). The single- and double-substitution substrates also substantially increase wild-type SmcR binding affinity for site E (Fig. 2B, 2E, and Table 1).

An interesting characteristic of the LuxR-family degenerate consensus sequence is that it is comprised of a combination of activated and repressed sites (23). It has been previously published that analysis of binding sites in promoters repressed by LuxR generates a consensus binding site with dyad symmetry with inverted repeats (Fig. 2C) (23). Conversely, sites associated with activated genes generates a consensus site with asymmetry; only one side of the palindrome is conserved (Fig. 2D). Thus, the combination of activated and repressed sites is what produces an asymmetric palindrome motif (Fig. 2B). The sequences recognized by SmcR N55I, including the P_*05222*_ site and the modified P_*luxC*_ site E, closely resemble the consensus for repressed binding sites (Fig. 2C). Conversely, SmcR N55I does not bind to sites that deviate from the repressed site consensus, such as most other sites in the *luxC* promoter (Fig. S2B). From these data, we conclude that SmcR N55I binds to a limited and specific subset of DNA sequences.

### SmcR N55I has reduced DNA binding and transcription activation at the *vvpE* promoter in *V. vulnificus*

We next wanted to study the relevance of the SmcR N55I substitution within the context of *V. vulnificus.* To examine the effects of the N55I substitution on SmcR transcriptional regulation *in vitro* and *in vivo* in *V. vulnificus*, we first used EMSAs to examine DNA binding at promoters in the SmcR quorum-sensing regulon. Two well-studied promoters from *V. vulnificus* were selected: P_*vvpE*_, which drives expression of elastase and is activated by SmcR, and P_*vvpM*_, which drives expression of a metalloprotease and is repressed by SmcR (31, 32). Purified SmcR N55I binds to the *vvpM* promoter with a slightly decreased affinity compared to wild-type SmcR, whereas SmcR N55I shows substantially decreased binding to the *vvpE* promoter compared to wild-type (Fig. 3A). Alignment of the *vvpE* and *vvpM* binding sites to the consensus sequences shows that they most closely resemble the activated and repressed consensuses, respectively (Fig. 2C, 2D). These results support our conclusion that SmcR N55I functions at repressed promoter sites but is non-functional for binding at some activated promoter sites.

**Figure 3.**
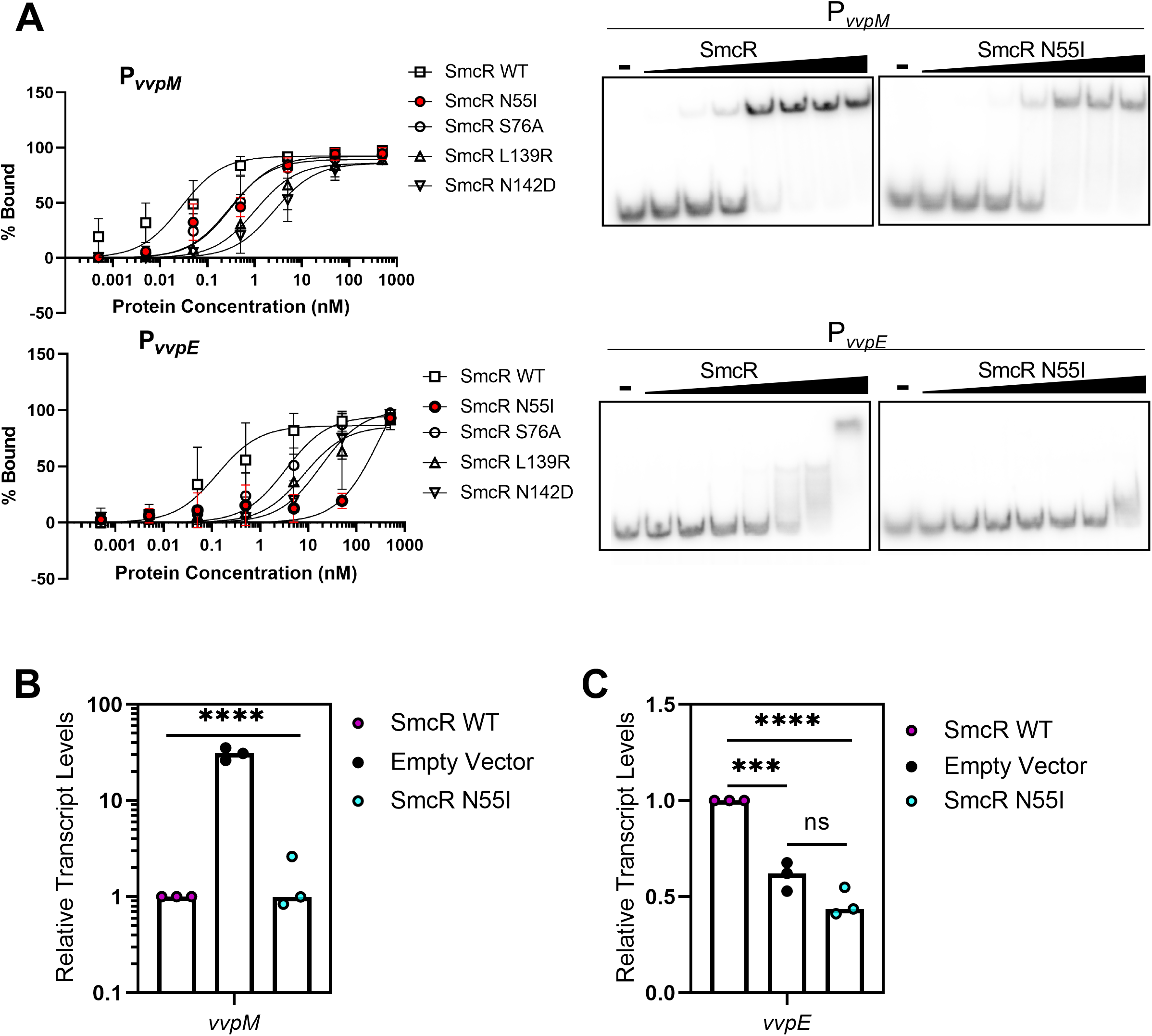
SmcR mutant N55I has limited DNA binding activity and transcriptional activation activity at P_*vvpE*_. **(A)** EMSA reactions consisting of 0.5 nM radiolabeled DNA substrates (P_*vvpM*_ or P_*vvpE*_ binding sites on the top and bottom, respectively, with oligonucleotides listed in Table 1 and S3) and increasing concentrations of purified wild-type SmcR or SmcR substitution mutant proteins (0.0005, 0.005, 0.05, 0.5, 5, 50, 500 nM protein). Data points show the average of three independent experiments, and error bars indicate the standard deviation. Representative gels of EMSA reactions are shown to the right. Lanes labeled ‘**—**’ had no protein added. **(B, C)** Relative transcript levels of *V. vulnificus* (B) *vvpM* and (C) *vvpE* determined by qRT-PCR of transcripts from *V. vulnificus* Δ*smcR* strains containing either a plasmid expressing *smcR* from an IPTG-inducible promoter (pJN22), *smcR* N55I (pJN27), or empty vector (pMMB67EH-kanR), induced with 50 μM IPTG. Asterisk (*) indicate that the fluorescence values are significantly different than the wild-type counterpart (**** *p* < 0.0001; *** *p* = 0.0005; one-way analysis of variance (ANOVA), followed by Tukey’s multiple-comparison test; *n* = 3).

We next assayed for transcription activation and repression activity at these promoters *in vivo* using qRT-PCR. The *smcR* or *smcR* N55I genes were expressed in a Δ*smcR* strain on an exogenous plasmid under control of a P_*tac*_ promoter, thus expression is induced by IPTG. Upon addition of 50 μM IPTG, expression of SmcR or SmcR N55I completely represses *vvpM* (Fig. 3B). However, while wild-type SmcR activates *vvpE* compared to the empty vector control strain, SmcR N55I does not increase *vvpE* expression, suggesting that it is not capable of activating the *vvpE* promoter (Fig. 3C). These data further support our conclusion that SmcR N55I is defective at binding specific DNA binding sites.

### The DNA-binding domain of wild-type SmcR exists in at least two conformations

We hypothesized that differences in the phenotypes observed between wild-type SmcR and the SmcR N55I proteins are due to structural differences in the DNA binding domains. We also predicted that any conformational changes would be subtle because the N55I mutant retains the ability to bind to a small subset of SmcR binding sites (Fig. 2A, S2A). In order to examine any differences between the structures, we first used X-ray crystallography to examine the structure of wild-type 6Xhis-tagged SmcR. The wild-type SmcR structure that we generated (6WAE; resolution 2.1 Å), as well as the published SmcR structure (3KZ9; resolution 2.1 Å) both have two dimers in the asymmetric unit (space group *P*2_1_2_1_2_1_; Fig. 4A) (33). Although our SmcR 6WAE structure contains a 19-residue N-terminal His-tag (for which no density is observed in the structure), it can be superimposed with the untagged SmcR structure with an average distance between alpha carbons (Cα) of 0.2 Å (Fig. S3). Throughout this manuscript, we refer to our His-tagged protein simply as SmcR and the previously solved untagged structure as SmcR 3KZ9. To quantify any differences between the two SmcR structures, we plotted the distances between alpha carbons of the four monomers for each of the two structures: SmcR A to SmcR 3KZ9 A, SmcR B to SmcR 3KZ9 B, SmcR C to SmcR 3KZ9 C, and SmcR D to SmcR 3KZ9 D (Fig. S3B, S3C). No major variations are observed in any of the four monomers alignments, with the exception of unstructured regions (*e.g.*, the N- and C-termini, Fig. S3C). We conclude from these data that the presence of the His-tag in the wild-type SmcR structure that we generated does not significantly affect the secondary structure.

**Figure 4.**
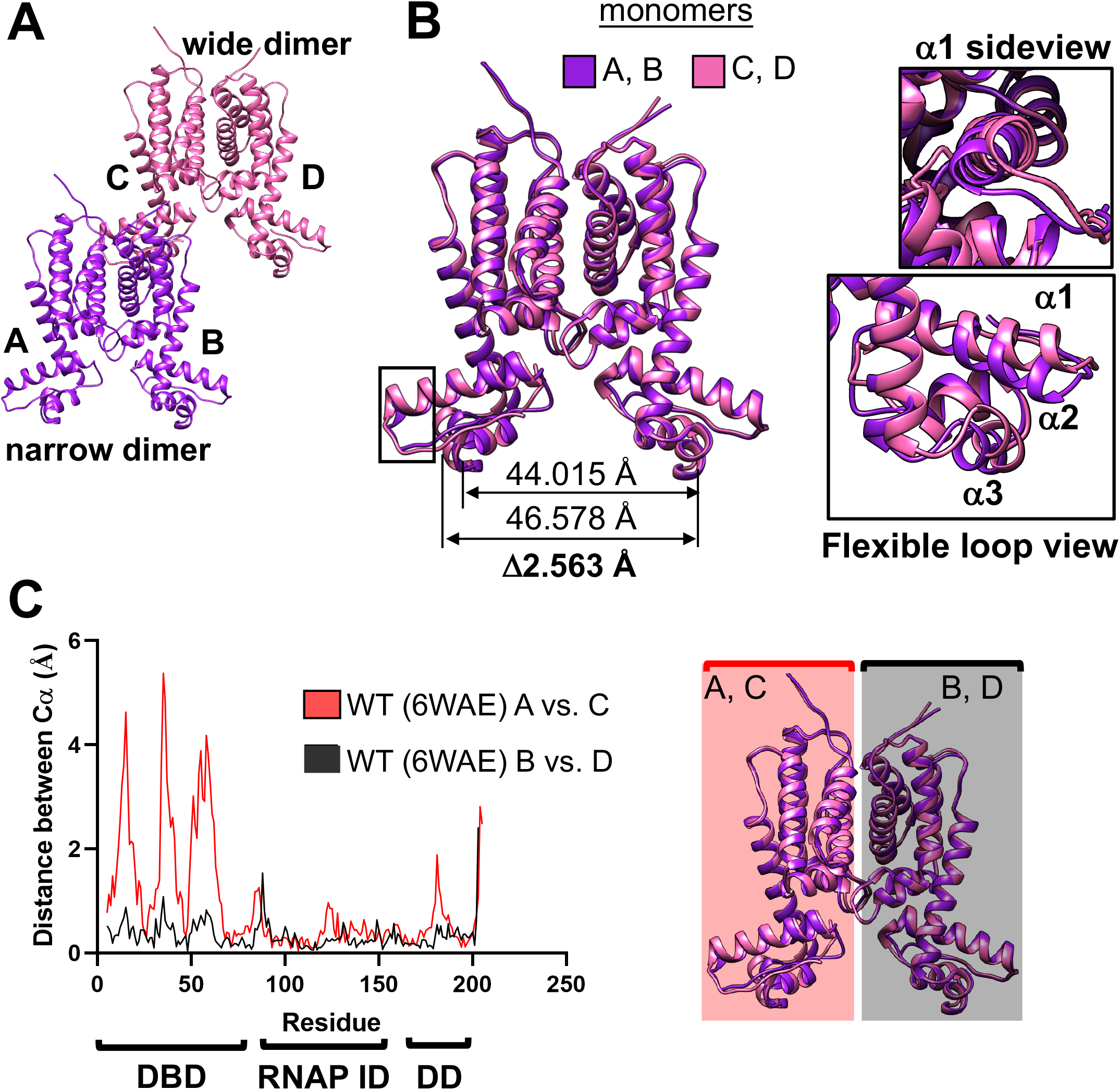
Wild-type SmcR has two DNA binding domain conformations. **(A)** The asymmetric unit containing two molecules of the wild-type His-tagged SmcR dimer (6WAE; purple and pink). **(B)** Crystal structure of the his-tagged SmcR wild-type (PDB: 6WAE) monomers A and B that form the narrow dimer (purple) are superimposed to monomers C and D that form the wide dimer (pink). The distance between alpha carbons (Cα) at residue 55 between monomers A to B (narrow dimer) or between monomers C to D (wide dimer) is shown beneath the structure (A-B measurement above C-D measurement). Helices 1, 2, and 3 are shown in insets to the right. **(C)** Graph showing the Cα of his-tagged SmcR wild-type (PDB: 6WAE) between monomer A to monomer C (red) and between monomer B to monomer D (black), with a diagram shaded in red or black to show which dimers are compared in the Cα graph. DBD, DNA binding domain; RNAP ID, RNAP interaction domain; DD, dimerization domain.

We next aligned the two SmcR dimers from the same asymmetric subunit by aligning the AB dimer to the CD dimer. The two dimers are remarkably similar except for two alpha helices and a loop in the DNA binding domain of one monomer that have a distinct shift (Fig. 4C). This shift exists in both wild-type SmcR structures: the His-tagged structure 6WAE and the previously solved SmcR 3KZ9 (Fig. S4A, S4B). We observe that the distances between DNA binding domain residues in the different dimers vary. For example, the distance between residue 55 in chain A and residue 55 in chain B of the AB dimer is 44.015 Å; whereas the distance between the same residues in the CD dimer is 46.578 Å (Fig. 4B). While this 2.6 Å change is not the largest observed (Fig. 4C), it is representative of the changes seen along the alpha helices in the DNA-binding domain. As a result, we refer to these two conformations as “narrow” (exhibited by SmcR dimer AB) and “wide” (exhibited by SmcR dimer CD). The individual Cα distances between the wide and narrow dimers in the DNA binding domain range from 2 to 6 Å approximately, whereas the average overall distances between individual Cα for comparing the wild-type wide and narrow dimers is 0.6 Å (Fig. 4C). It is important to note that in all crystallographic studies a caveat is that conformational changes may be artifacts due to crystal packing, so we have made a note of all crystal contacts in a figure (Fig. S7). Although we observe two conformations, we cannot rule out the possibility that the DNA binding domain dwells in additional conformations than the two captured in the crystal structures. Additional conformations could be intermediate states between the two conformations shown here or could be conformations sampling wider or narrower states than our solved structures.

### The N55I substitution limits the DNA-binding domain of SmcR to a single conformation

We also used X-ray crystallography to examine the structure of SmcR N55I and determined the structure at 3.4 Å. Unfortunately, our attempts to improve the resolution, including the crystallization of His-tagged variants, were unsuccessful. We next superimposed the structure of SmcR N55I (6WAF) to wild-type SmcR 3KZ9 since both structures were generated from untagged proteins (33). These superimpositions were performed using the matchmaker function in Chimera, which considers the best alignment for the overall structure (Fig. 5A). The algorithm can also consider the dimerization domains (the part of the protein suspected to have the least flexibility and lowest B-factor values) as the fixed region. However, we opted to utilize the matchmaker function in Chimera to analyze the data using the best overall fit to ensure less user bias. SmcR N55I has only one dimer in the asymmetric unit (6WAF; space group C2). This dimer aligns best to the wide dimer of wild-type SmcR 3KZ9 and seems to be distinct from the SmcR 3KZ9 narrow dimer (Fig. 5A). We quantified the structural differences by comparing the distances between alpha carbons in the SmcR N55I wide dimer to the SmcR 3KZ9 narrow dimer (Fig. 5B, 5C) or SmcR 3KZ9 wide dimer (Fig. 5D, 5E). The alignment of SmcR N55I to the SmcR 3KZ9 wide dimer plot shows little variation, with distances between alpha carbons generally less than 2 Å (Fig. 5D, 5E). However, the SmcR N55I monomer A shows large (2 - 4 Å) variations in Cα compared to SmcR 3KZ9 monomer A in the narrow dimer in the DNA-binding domain residues 5 to 75 (Fig. 5B), reminiscent of the variations observed between the wide and narrow dimers in SmcR wild-type (Fig. 4C). The fluctuations in the other regions of the proteins are small (protein’s overall Cα average = 0.76 Å), especially in flexible loop regions as expected (Fig. 5B-E). It is noteworthy to mention that despite the low resolution at which SmcR N55I was solved, alpha-carbon distances estimated in this work are rather accurate. Hardly surprising, the side chains in low resolution structures are often mistakenly assigned due to the deficient electro-density maps in those regions. However, generally the electron-density for the protein backbone, as well as the alpha-carbon atoms assigned are more certain. Because the SmcR N55I structure was solved at a lower resolution, the B-factors are higher values compared to the SmcR wild-type structure (Table S4). However, when considering the overall range of B-factors in the SmcR N55I structure, the relative values for the DNA binding domain compared to the dimerization domain are similar to that of wild-type (33). Additionally, any shifts noticed in the DNA-binding domain do not appear to be directly due to crystal contacts, as these are mostly apparent near the top of the dimerization domain and the flexible loops of the protein (Fig. S7). There are many crystal contacts displayed in the N55I structure and wild-type structure so we cannot rule out the influence of crystal contacts on conformational changes. While we cannot solely rely on crystallography to assess these conformational changes, our model of the DNA-binding domain adopting multiple conformations is not new. Another group has used small/wide angle X-ray scattering (SWAXS) to observe a similar phenomenon in HapR (34). As this technique is performed using protein in solution, rather than crystallized protein, we have increased confidence that these conformational changes are not due to crystal packing artifacts. Together, these results indicate that the single SmcR N55I structure aligns closely with one conformation of wild-type SmcR (the wide dimer) but not the other (narrow dimer).

**Figure 5.**
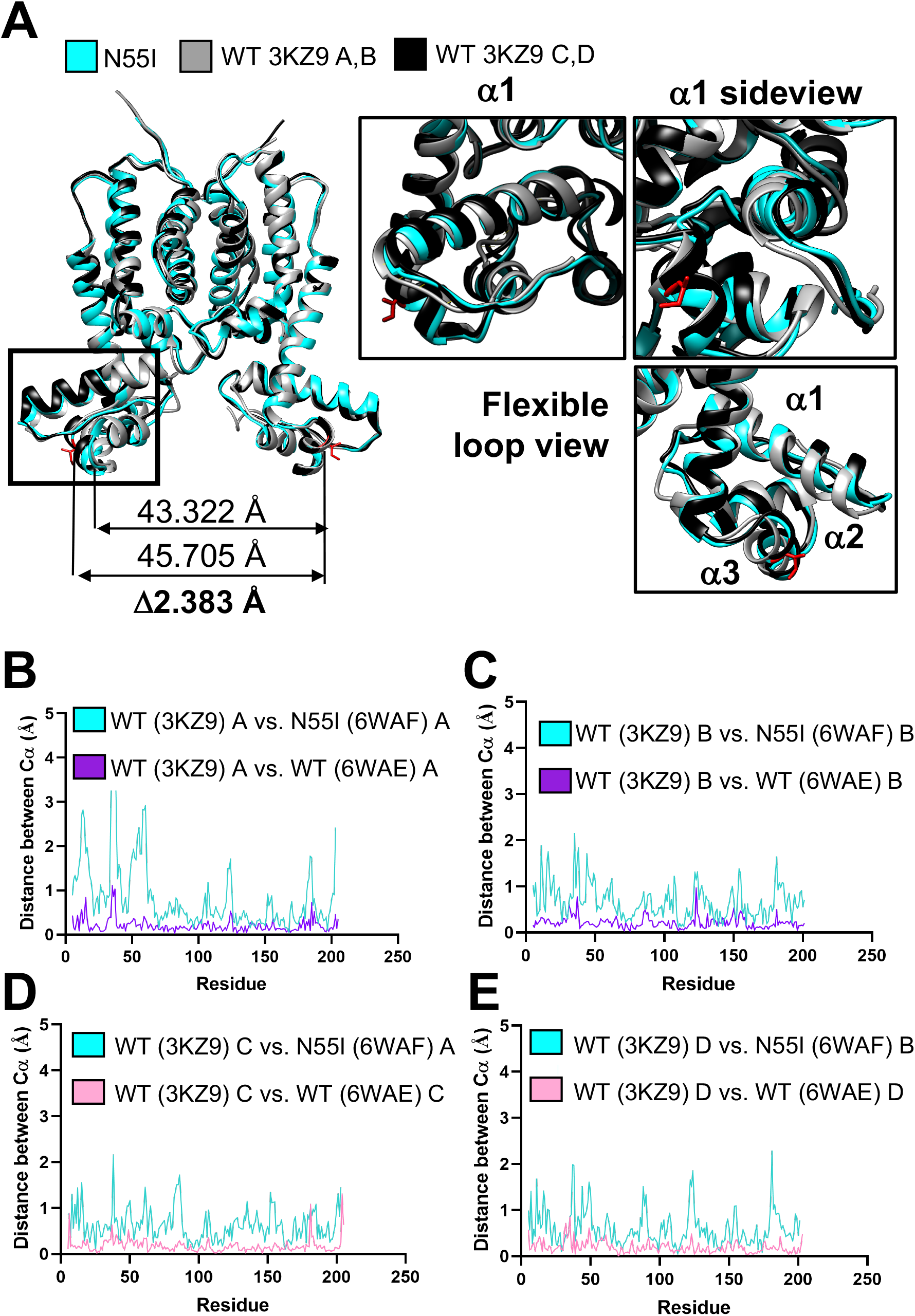
Crystal structure of SmcR N55I exists in the wide conformation. **(A)** The two dimers of SmcR 3KZ9 (narrow AB dimer, gray; wide CD dimer, black) and SmcR N55I dimer (cyan; 6WAF) superimposed. The insets show the superimposed α1, α2, and α3 helices and the connecting loop in the DNA binding domain. The distance between alpha carbons (Cα) at residue 55 between SmcR 3KZ9 monomers A to B (narrow dimer) or between SmcR N55I A to B (similar to WT C and D; wide dimer) is shown beneath the structure (3KZ9 measurement above N55I measurement). **(B, C, D, E)** Graphs of the Cα distances calculated by Chimera. Compared distances are as follows: **(B)** SmcR wild-type monomer A versus SmcR N55I monomer A (cyan), and SmcR 3KZ9 monomer A versus SmcR 6WAE monomer A (purple), **(C)** SmcR 3KZ9 wild-type monomer B versus SmcR N55I monomer B (cyan), and SmcR 3KZ9 monomer B versus SmcR 6WAE monomer B (purple), **(D)** SmcR wild-type monomer C versus SmcR N55I monomer A (cyan), and SmcR 3KZ9 monomer C versus SmcR 6WAE monomer C (pink), and **(E)** SmcR wild-type monomer D versus SmcR N55I monomer B (cyan), and SmcR 3KZ9 monomer D versus SmcR 6WAE monomer D (pink).

### SmcR RNAP-interaction mutants have two-conformation structures similar to wild-type

The amino acid substitutions S76A, L139R, and N142D in SmcR alpha helices 4 and 7 are predicted to be involved in protein-protein interactions based on previous research on LuxR and HapR, given that these substitutions render the proteins unable to interact with RNAP alpha *in vitro* and are decreased in transcription activation (28). To determine whether these substitutions affect the structure of the protein, we solved the X-ray crystal structures of SmcR S76A, L139R, and N142D (resolutions of 2.57 Å, 2.55 Å, and 2.58 Å, respectively; PDB 6WAG, 6WAH, 6WAI). Each of the substitution mutants has two dimers in the asymmetric unit. These structures are highly similar to wild-type SmcR and show no appreciable change in alpha helices from the wild-type SmcR structure aside from movement in unstructured loops. The average overall alpha carbon distance between wild-type and the substitution mutants S76A, L139R, and N142D are 0.18 Å, 0.25 Å, and 0.17 Å, respectively (Fig. S5). Importantly, much like wild-type SmcR, each of the substitution mutants also show two DNA binding domain conformations (Fig. 6A, 6C, 6E). The alpha carbon distances between the wide and narrow dimers for each substitution mutant show that the DNA binding domain alpha helices are also shifted, similar to wild-type SmcR (Fig. 6B, 6D, 6F). From these structural data, we conclude that the S76A, L139R, and N142D substitution mutants of SmcR have similar structures to wild-type SmcR and exist in multiple conformations.

**Figure 6.**
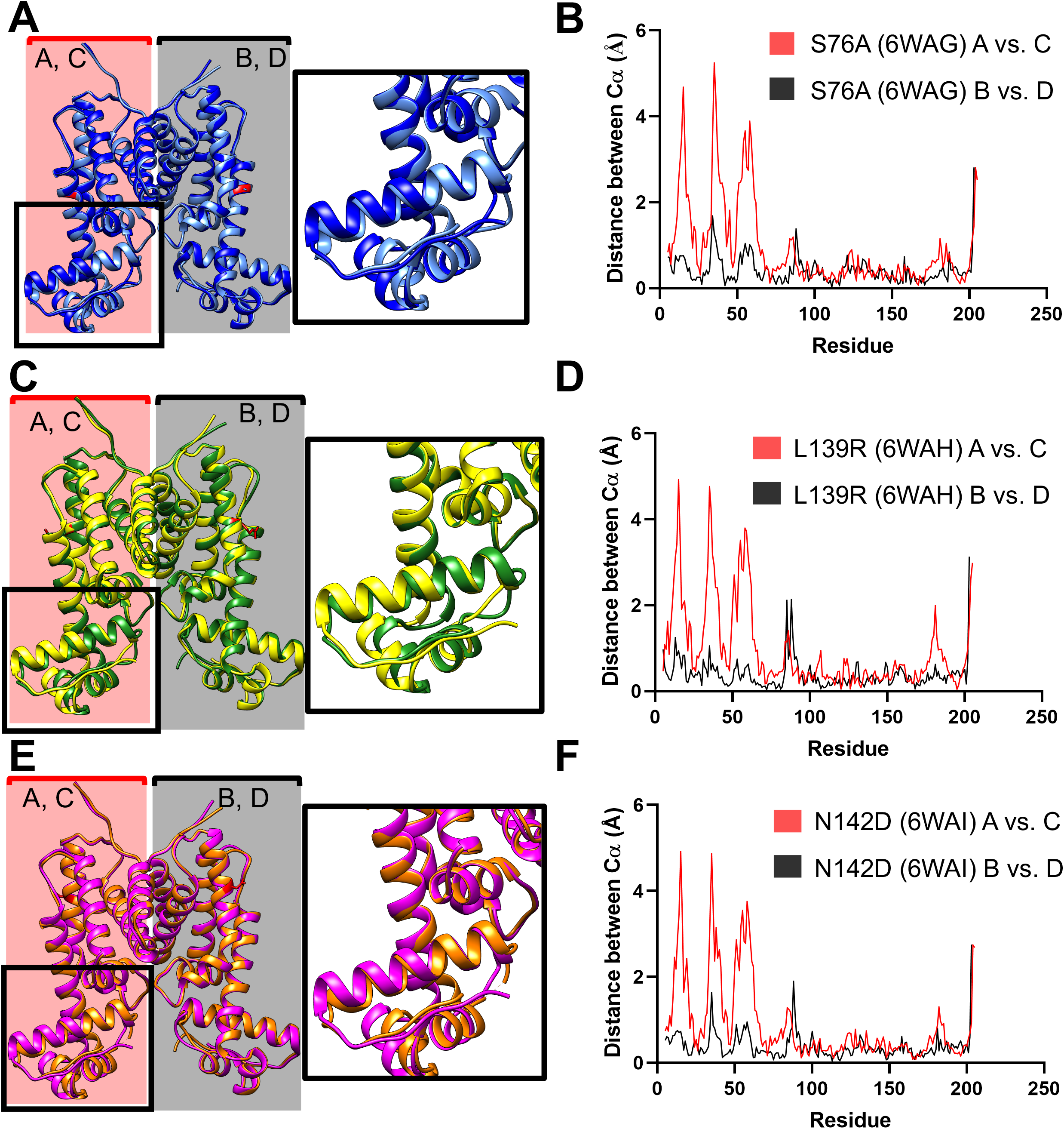
SmcR S76A, L139R, and N142D exhibit structures similar to wild-type with two DNA binding domain conformations. **(A)** SmcR S76A (6WAG) narrow dimer (cornflower blue) structure aligned to the wide dimer (royal blue) with an inset zoomed into one monomer of the DNA binding domain. **(B)** Graph showing the Chimera measured distances between alpha carbons (Cα) of his-tagged SmcR S76A (PDB: 6WAG) from monomer A to monomer C (red) and from monomer B to monomer D (black). **(C)** SmcR L139R (6WAH) narrow dimer (forest green) structure aligned to the wide dimer (yellow) with an inset zoomed into one monomer of the DNA binding domain. **(D)** Graph showing the Cα of his-tagged SmcR L139R (PDB: 6WAH) from monomer A to monomer C (red) and from monomer B to monomer D (black). **(E)** SmcR N142D (6WAI) narrow dimer (orange) structure aligned to the wide dimer (magenta) with an inset zoomed into one monomer of the DNA binding domain. **(F)** Graph showing the Cα of his-tagged SmcR N142D (PDB: 6WAI) from monomer A to monomer C (red) and from monomer B to monomer D (black).

Based on these data, we predicted that the DNA binding activity of these mutants is similar to wild-type SmcR. To test this, we used EMSAs to assay DNA binding at the *vvpE* and *vvpM* promoters for each substitution mutant compared to wild-type SmcR. The binding activities of L139R, N142D, and S76A SmcR mutant proteins are decreased compared to wild-type SmcR at both the *vvpE* and *vvpM* promoters (Fig. 3A, S6). The *K*_D_ of the RNAP-interaction substitution mutants with P_*vvpM*_ (S76A = 0.3 nM, L139R = 1 nM, N142D = 2.6 nM) and N55I (*K*_D_ = 0.3 nM) vary greatly from wild-type (WT = 0.03 nM), but these variations are most likely not physiologically relevant given that these are all very high affinities at ~3 nM or less. The decreases in affinity for P_*vvpE*_ and the RNAP-interaction mutants are larger (S76A = 3.4 nM, L139R = 8.2 nM, N142D = 18.3 nM) but still well within the range of transcription factor binding affinities in other bacterial regulatory systems. However, the N55I *K*_D_ of 303.5 nM is comparatively a very poor binding affinity for DNA (19, 35, 36). It is possible that the decreased DNA binding affinity of the RNAP-interaction mutants may contribute to the decreased transcription activation of these mutant proteins *in vivo*. However, this modest decrease in DNA binding ability does not affect the N-terminal DNA-binding domain conformations, and thus appears to be different from the N55I conformation and associated DNA-binding defects.

## Discussion

The study of transcriptional control of quorum-sensing genes is pivotal for a basic biological understanding of the evolution of coordinated behaviors, as well as for a better understanding of key drug targets to treat diseases coordinated by quorum sensing. Notably, LuxR/HapR/SmcR proteins are members of the TetR family of transcription factors, yet they have evolved to bind more than 100 sites throughout the genome. This study has investigated how these regulators simultaneously maintain specificity for quorum-sensing targets and diversity to accommodate binding at numerous promoters for both activation and repression activities.

Ultimately, information regarding the mechanism of transcriptional regulation of the LuxR/HapR/SmcR family of proteins will inform future studies aiming to disrupt these pathways to mitigate pathogenesis regulated by quorum sensing (37, 38). Specifically, the protein-protein and protein-DNA interactions required for function in these regulators represents a key target for small molecule inhibitors. Numerous small molecule inhibitor studies have been carried out targeting QacR, a homologue from *Staphylococcus aureus*, which also happens to be one of the few LuxR/HapR/SmcR family of proteins to be co-crystallized bound to DNA (Fig. 7A). By comparing the QacR-DNA structure to QacR-drug bound structures, we observed shifts in the DNA-binding domain with limited motion in the dimerization domain. While the shift in the DNA-binding domain is distinct from the shifts observed in this study, we anticipate that conformational flexibility in the DNA-binding domain allows for substrate binding at higher affinities with increased specificity. QacR bound to molecules that inhibit QacR function have a wider conformation than the DNA-bound QacR, which supports our model that wide conformations have a more limited function *in vivo* (Fig. 8). While none of the *Vibrio* proteins have been successfully co-crystallized with DNA, the QacR structure does give insights into putative structural shifts occurring in these proteins upon DNA binding. We hypothesize that upon SmcR binding DNA, a more substantial shift may likely exist, however our extensive trials with co-crystalization have thus far been fruitless. Similar to the *S. aureus* inhibitor, studies in vibrios have produced some small molecule inhibitors with promising inhibitory effects on LuxR/HapR/SmcR, such as Qstatin, which limits transcription regulation by SmcR *in vivo* (15, 39, 40). The Qstatin-bound SmcR structure also exhibits the same two conformations in the DNA binding domain, suggesting that the mechanism of Qstatin inhibition is not due to limitation of DNA binding sequence recognition. Further mechanistic studies will be required to determine the way by which Qstatin inhibits SmcR function *in vivo*.

**Figure 7.**
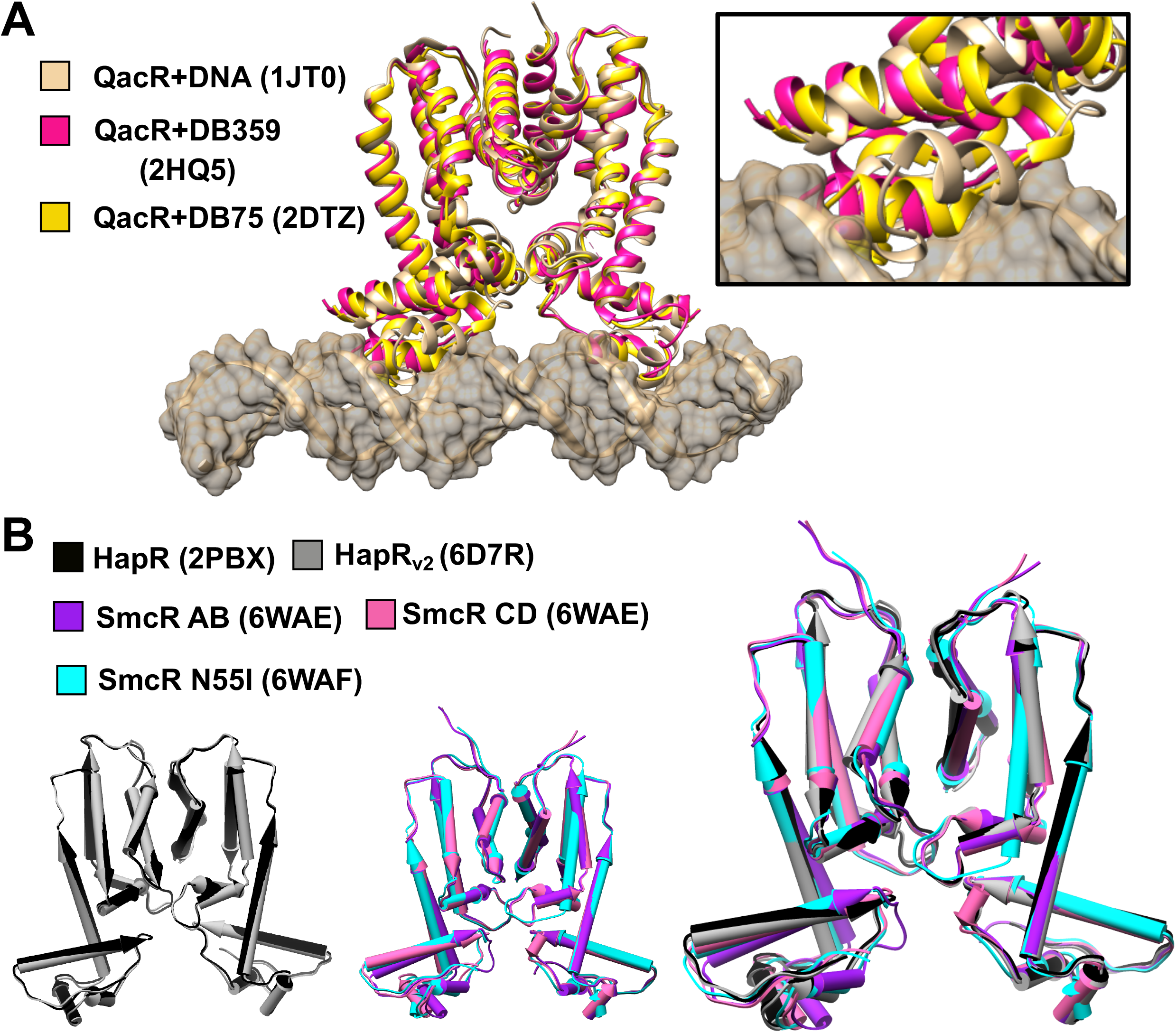
Structural comparisons of homologous TetR family proteins. **(A)** Structure of QacR and DNA (tan) superimposed with QacR bound to inhibitors DB359 (pink) and DB75 (gold). Inset shows zoomed in view of DNA-binding domain. **(B)** (left) Structure of HapR wild-type (black) superimposed with HapR inactive variant (HapR_V2_; gray). (middle) SmcR superimposition of wild-type dimers AB (purple), CD (pink), and SmcR N55I (cyan). (right) All HapR and SmcR superimpositions.

**Figure 8.**
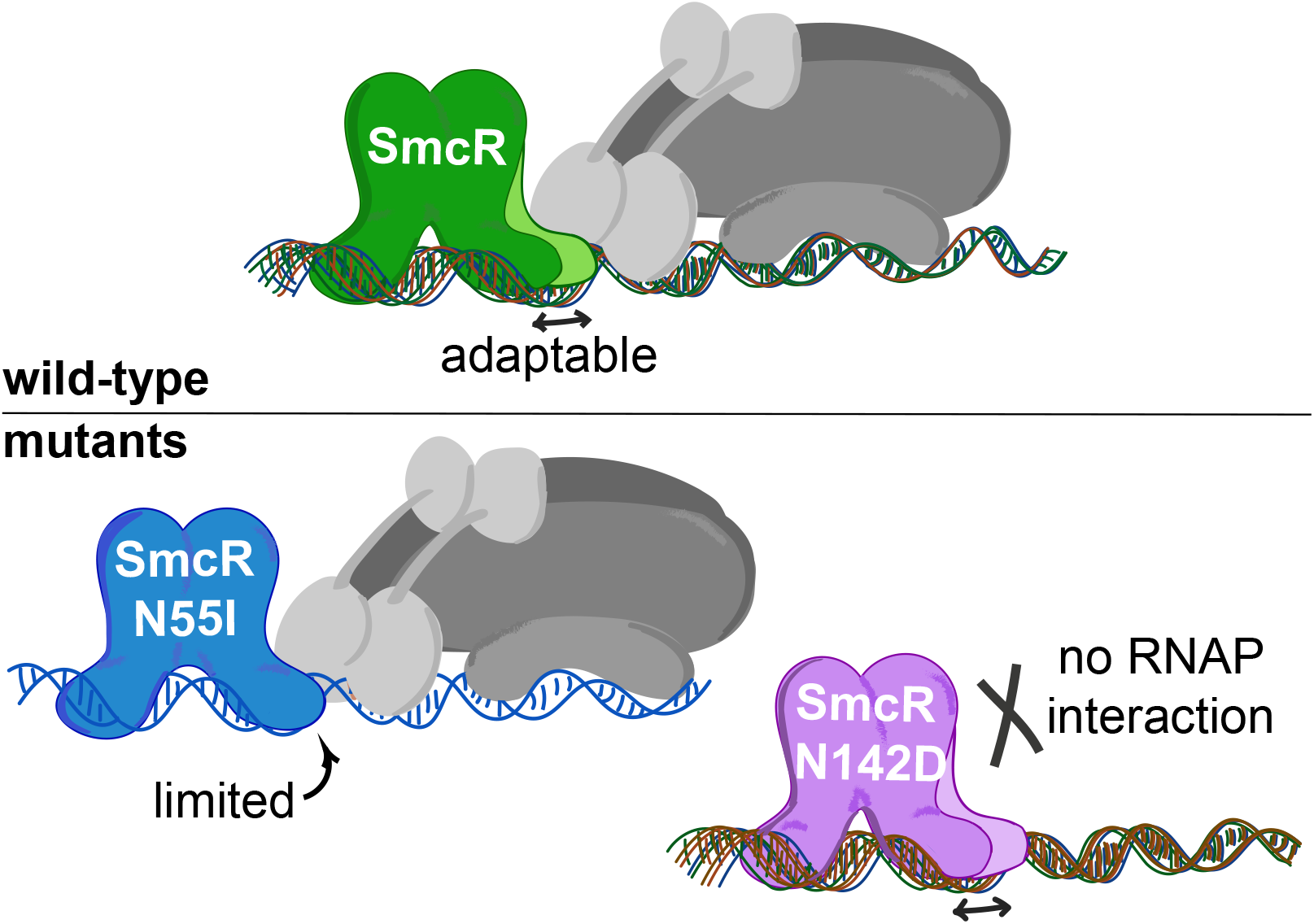
Model: Multiple SmcR DNA binding domain conformations accommodate a wide array of DNA substrates. One DNA binding domain in SmcR exhibits at least two conformations in crystal structures, enabling recognition of numerous DNA sequences. Mutant proteins have loss-of-function transcriptional phenotypes due to either 1) a single DNA binding domain conformation conferring limited sequence recognition (*e.g.*, N55I), or 2) disrupted interaction with RNAP but maintained DNA binding activity (*e.g.*, N142D).

Interestingly, the wild-type HapR crystal structure (PDB: 2PBX) and an inactive DNA-binding HapR naturally occurring variant (HapR_V2_; PDB: 6D7R) both only show one conformation in their crystal structures, which align closest to the wide SmcR conformation (Fig. 7B)(41). Importantly, the HapR crystal structures are in the same space group as the wild-type SmcR structures, so we compared the crystal packing (Fig. S7). The crystal contacts observed for HapR and HapR_V2_ vary significantly, yet the conformations align very closely, also suggesting that the contacts play a minimal role in the conformations present. This further supports the idea that N-terminal domain conformation adaptability is only one mechanism utilized by LuxR/SmcR/HapR, in addition to interactions with RNAP and DNA (28). It is interesting to consider whether the single conformation observed in the HapR structures indicates that it has a limited DNA binding consensus. To our knowledge, the available HapR binding site consensus sequence was generated primarily through bioinformatics analysis of a handful of known regulated sites (22), but global ChIP-seq studies or similar experiments have not been performed to examine the sequences directly bound by HapR *in vivo*. Further studies of the HapR DNA binding site sequences will be informative to determine if HapR possesses fewer or more constrained conformations to restrict DNA sequence recognition

Our structural studies have shown that multiple structural conformations are adopted by SmcR to facilitate binding of variable DNA substrates in quorum-sensing promoters. The measurable alteration in the SmcR dimer between the two helix-turn-helix domains may enable SmcR to interact with DNA sequences from various promoters where sequence variation is restricted. For example, many LuxR binding sites have been shown to overlay the −35 region bound by sigma-70 (23, 24, 30). Limitations to changes in these regions may have driven selection for the protein to adapt to variable sequences, rather than mutations to the binding site itself. Importantly, the DNA-binding defective SmcR N55I mutant provided the key needed to understand the function of these multiple conformations. The N55I protein, with only one conformation in the crystal structure, has a limited range of sequence recognition. Conversely, wild-type SmcR binds to a wide range of DNA sequences, resulting in the degenerate asymmetric binding consensus for this family of proteins. Thus, we propose that the mechanism by which SmcR recognizes hundreds of binding sites with variable sequences is by employing a DNA binding domain that is adaptable to accommodate diverse sequences (Fig. 8). The SmcR N55I protein has a more limited flexibility and therefore more limited range of DNA sequences to bind. We note that the sequences bound by SmcR N55I represent sites at which LuxR has been shown to have the tightest binding affinity (*e.g.*, P_*luxC*_ site H and P_*05222*_) (23). This could indicate that LuxR/HapR/SmcR proteins have evolved to be increasingly flexible to accommodate more binding sites for an expanded regulon compared to traditional TetR-type proteins. This hypothesis could be addressed with future evolution experiments evolving a canonical TetR-type protein to bind to additional promoters, such as the bioluminescence promoter P_*luxC*_.

The focus of this study has been solely on the biophysical mechanisms of activation and how specific SmcR-DNA and SmcR-protein interactions influence SmcR regulation of activated genes. We took advantage of known activation-deficient mutants to determine how these substitutions affected SmcR activity both *in vitro* and *in vivo*. Similar analyses could be performed to interrogate repression-deficient mutants to study whether similar SmcR-DNA interactions are vital for SmcR to function as a repressor (23). One challenge we have already faced with the repression-deficient substitution mutants is the inability to crystalize these proteins using the same purification and crystallization schemes we used for the proteins presented in this manuscript. This may indicate differences in solubility or flexibility of these proteins, and perhaps other techniques such as NMR would be beneficial to further understand the structural differences in these proteins. The DNA-bound SmcR structure would truly contribute the greatest information in DNA binding site recognition, though numerous attempts with various DNA substrates, protein purification schemes, and a library of crystallization conditions has not yet yielded crystals that diffract.

The core importance of the work presented here is that the *Vibrio* family of LuxR/HapR/SmcR proteins have evolved from traditional TetR-type proteins and have the ability to activate transcription and to bind to numerous and diverse DNA sequences. Our model is that two biophysical properties enable these activities: 1) interaction with RNAP, which likely recruits or stabilizes RNAP at certain promoters, and 2) multiple DNA binding domain conformations that allow recognition of various sequences (Fig. 8). While the hypothesis that LuxR/HapR/SmcR proteins adopt multiple conformations is not new, the biochemical, structural, and quantitative protein comparisons data presented in this manuscript are the first to show definitive support for this model.

## Materials and Methods

### Bacterial strains and media

*E. coli* strains DH10B and S17-1λpir were used for cloning, and BL21(DE3) was used for overexpression of proteins. All *E. coli* strains and derivatives were grown in Lysogeny Broth (LB) at 30°C shaking at 275 RPM in LB media with the corresponding antibiotic (Table S1). *V. vulnificus* ATCC 27562 and derivatives were grown shaking at 275 RPM at 30°C in Luria Marine (LM) medium (LB with 2% NaCl) with appropriate antibiotics. Antibiotics were used at the following concentrations: kanamycin 50 μg/mL or 250 μg/mL (*E. coli* or *V. vulnificus*, respectively), chloramphenicol 10 μg/mL, ampicillin 100 μg/mL, and tetracycline 10 μg/mL. The dual promoter fluorescence reporter assays were performed using plasmid pJV064 containing the P_*luxC*_ fused to GFP and P_*05222*_ fused to mCherry to assess LuxR transcriptional regulation. Overnight *E. coli* cultures containing *luxR* and the dual fluorescence reporter were diluted 1:1000, induced with 15.6 nM IPTG, and grown for 16 hours at 30°C. The OD_600_ and fluorescence (both GFP and mCherry) was measured on a BioTek plate reader.

### Molecular methods

PCR was performed using Phusion HF polymerase purchased from New England Biolabs (NEB). T4 polynucleotide kinase (T4 PNK) used in EMSAs and all other enzymes mentioned were purchased from NEB and used according to manufacturer’s instructions. Site-directed mutagenesis for overexpression of mutant proteins was carried out using the Agilent QuikChange II XL Site-Directed Mutagenesis Kit. The mutations were confirmed by DNA sequencing (Eurofins). All oligonucleotides were purchased from Integrated DNA Technologies (IDT), and those used in this study are listed (Table S3). Cloning details for plasmids listed in Table S2 are available upon request.

### RNA analysis by qRT-PCR

To collect RNA samples, cells were grown at 30°C shaking at 275 RPM to an OD_600_ of approximately 0.2. Then cells were induced with 50 μM IPTG and grown under the same conditions until cells reached an OD_600_ of approximately 1, at which 5 mL of cells were collected by centrifugation and frozen in liquid N_2_. RNA was extracted using a Trizol/chloroform extraction protocol previously described (43) and cleaned up using an RNeasy Mini Kit (Qiagen).

Quantitative real-time PCR (qRT-PCR) was performed as previously described (30). Samples were normalized to the internal standard *recA* gene. The ΔΔ*C*_*T*_ values were used to analyze data from three independent biological replicates. Symbols on graphs represent the mean values and error bars represent the standard deviations. All statistical analysis was performed with functions from GraphPad Prism version 8. Further details are available in the figure legends.

### Purification of proteins

SmcR and RNAP alpha proteins were purified similarly to previous publications with minor adjustments (23, 28, 44). *E. coli* BL21(DE3) strains containing plasmids expressing hexahistidine-tagged *smcR* wild-type and mutant alleles were grown overnight in LB medium with kanamycin, back-diluted 1:100 into 1 L of LB medium with kanamycin, and grown to an OD_600_ of 0.4-0.6 at 30°C. Expression of SmcR was induced by IPTG to a final concentration of 1 mM and cultures grown for 4 h shaking at 30°C. The cells were pelleted and frozen at −80°C. The pellet was resuspended in 25 mL buffer A (25 mM Tris pH 8, 500 mM NaCl), and an Avestin EmulfiFlex-C3 emulsifier was used to lyse cells. The soluble lysate was applied to a HisTrap HP Ni-NTA column using an Äkta Pure FPLC in buffer A and eluted from the column with a gradient of buffer B (25 mM Tris pH 8, 500 mM NaCl, 1 M imidazole). The purified protein was concentrated to approximately 5 ml using Sartorius Vivaspin Turbo 10,000 MWCO centrifugal concentrators. The sample was manually injected into the Äkta Pure and separated via size exclusion chromatography on a HiLoad™ 16/600 Superdex™ 75 pg column equilibrated with gel filtration buffer (25 mM Tris pH 7.5, 200 mM NaCl). Eluted fractions were analyzed by SDS-PAGE, pooled, and concentrated using the same centrifugal concentrators previously mentioned. The samples were then immediately used in crystal trays or frozen in liquid nitrogen with a final concentration of 10% glycerol and stored at −80°C. All SmcR proteins used in Bio-layer Interferometry BLI) experiments and SmcR N55I for crystallography was overexpressed in *E. coli* BL21(DE3) using the Intein Mediated Purification with an Affinity Chitin-binding Tag (IMPACT) system utilizing the pTXB1 vector for a C-terminal tag. Cells were grown and induced the same as previously stated, pelleted, and resuspended in buffer 1 (25 mM Tris pH 8, 500 mM NaCl, 1 mM EDTA) prior to lysis by an Avestin EmulfiFlex-C3 emulsifier. The soluble lysate was applied to NEB chitin resin in buffer 1. The resin was washed with 10 CV buffer 1 and 10 CV buffer 2 (25 mM Tris pH 8, 1 M NaCl, 1 mM EDTA) then incubated with 30 mL buffer 1 + 231 mg dithiothreitol (DTT) for 4 hours to promote self-cleavage of the intein tag from the SmcR proteins. The protein cleaved from the intein-tag was eluted with buffer 1 and dialyzed for 2 hours in gel filtration buffer. The dialyzed fractions were then concentrated to ~3 mL and manually injected into the Äkta Pure and separated via size exclusion chromatography on a HiLoad™ 16/600 Superdex™ 75 pg column equilibrated with gel filtration buffer. The purification of his-tagged *V.harveyi* RNAP α was previously described (28).

### Crystallization and structure determination of SmcR variants

SmcR wild-type and variants were crystallized in conditions similar to those previously described (33) with slight variations. SmcR WT and variants (~4 mg/mL) crystals grew at 20°C using the hanging-drop vapor-diffusion method. Crystals formed under the condition of 0.2 M of lithium sulfate, 0.1 M imidazole buffer pH 7.6-8, and 6-10% PEG3350. Then, crystals were harvested, cryo-protected in reservoir solution supplemented with 20% ethylene glycol or a mix of 10% glycerol and 10% ethylene glycol and flash-frozen in liquid nitrogen. Diffraction data were collected at 100 K at the Beamline station 4.2.2 at the Advanced Light Source (Berkeley National Laboratory, CA) and were indexed, integrated, and scaled using XDS (45). The structures were solved by molecular replacement using PHASER and the PDB code 3KZ9 (only one chain) as a search model. The Autobuild function was used to generate a first model that was improved by iterative cycles of manual building in Coot (46) and refinement using PHENIX (47). MolProbity software (47) was used to assess the geometric quality of the models and Chimera (48) or Pymol (49) to generate molecular images. Data collection and refinement statistics are indicated in Table S4. All data sets but variant N55I (space group C2) were initially processed in space group *P*2_1_2_1_2_1_. Data presented translational pseudo-symmetry (as defined by Xtriage in Phenix), patterson peaks with length larger than 15 Å and pseudo-translation vector (0.234, 0.5, 0.0). As a result, multiple molecular replacement solutions with high TFZ and LLG were obtained, most of them with R-free values over 0.34 after refinement. Then, data sets were reprocessed in lower symmetry space groups. All solutions from molecular replacement, in every possible space group, were built and refined. Those with the best R factors were achieved in space group *P*2_1_2_1_2_1_ (Table 1). Translational non-crystallography correction was not used during refinement, since it had marginal effect, if any, on R values. UCSF Chimera software was utilized to superimpose structures using the matchmaker function and then distances between alpha carbons were calculated. These were calculated using the following command: distance #1.1 :5.a@CA #2.1 :5.a@CA. This comparison was done for each residue present in the solved structures. Chimera was also used to display crystal contacts shown in figures, as well as measure the distances of these putative contacts. This was done using the Tools: Higher-Order Structure function and selecting Multiscale Models. Then with loaded atoms and contact distance near a set range of 5 Å, select the Multimer 3×3×3 crystal unit cells and make models. Then clashes/contacts can be utilized from the structural analysis function in tools. The parameters used for contacts was the default criteria: VDW overlap = − 0.4 Å and subtract 0 from overlap for potentially hydrogen-bonding pairs. All protein structures in figures were produced using Chimera software.

### Radiolabeling DNA Probes and Electrophoretic Mobility Shift Assays (EMSAs)

The EMSAs were conducted in a similar fashion to previous publications (28, 30). The oligonucleotides used as substrates were annealed in 1X annealing buffer (50 mM Tris-HCl pH 7.5, 100 mM NaCl) at 95°C for 1 min then cooled from 95°C-10°C at 1°C/min to generate dsDNA substrates. The 500 nM annealed DNA substrates were then radiolabeled in PNK buffer with PNK enzyme and radioactive ATP [γ^32^P] from Perkin Elmer at 37°C for 1 hr then diluted in TE buffer (10 mM Tris, 1mM EDTA pH 8). The excess ATP [γ^32^P] was removed using GE Healthcare G-25 columns. The final dsDNA concentration of the probes was 5 nM. EMSAs were carried out as previously described (30) with a dilution series of protein (specified in figure legends), 10 ng/μL poly(dIdC), 100 μg/mL bovine serum albumin (BSA), 1X binding buffer (10 mM HEPES pH 7.5, 100 mM KCl, 2 mM DTT, 2 mM EDTA) for 30 mins at 30°C. Protein-DNA complexes were visualized on TGE (25 mM Tris, 250 mM glycine, 1 mM EDTA) polyacrylamide gels in TGE buffer. Gels were dried for 1 hour at 80°C then exposed to a phosphor screen and analyzed on a Typhoon 9210 (Amersham Biosciences).

### Bio-layer interferometry

Bio-layer interferometry (BLI) was performed according to protocols previously established (28). BLI was performed on an Octect® K2 System using Dip and Read™ Ni-NTA (NTA) Biosensors (FortéBio®). In a 96-well plate, the interaction of untagged SmcR proteins (WT or N55I) were assayed with 200 nM his-tagged *V. harveyi* RNAP α subunit. His-tagged Tobacco etch virus (TEV) protease was used as a control for non-specific interactions (28). The reactions were carried out in SmcR gel filtration buffer (25 mM Tris pH 7.5, 200 mM NaCl) with various concentrations (1000, 850, 700, 550, 400, 250, and 0 nM) of the SmcR protein (*i.e.,* the analyte). The method on the Octet® K2 System Data Acquisition 9.0 software was set up to perform 30 s sensor equilibration in the reference well, 300 s ligand-loading step in the ligand-loading well, 60 s baseline step in the reference well, 700 s association step in the association well, and 700 s dissociation step in the reference well. All steps were performed shaking at 30°C for each concentration of analyte. The data were analyzed using Octet® K2 System Data Analysis 9.0 software.

## Acknowledgements

The authors gratefully acknowledge use of the Macromolecular Crystallography Facility (MCF) in the Molecular and Cellular Biochemistry Department, Indiana University Bloomington. We also thank the Indiana Clinical and Translational Science Institute (CTSI) grant for Core Facility Funds supporting experimentation in the MCF. We also thank Jay Nix for his assistance during X-ray data collection at beamline 4.2.2, ALS. This project was funded by National Institutes of Health grant R35GM124698 to JVK.

## Author Contributions

JN and JVK designed the experiments, JN, MR, GGG, and JVK performed experiments, JN, GGG, and JVK analyzed results, and JN, GGG, and JVK wrote the manuscript.

## Supporting Information

**Figure S1.**
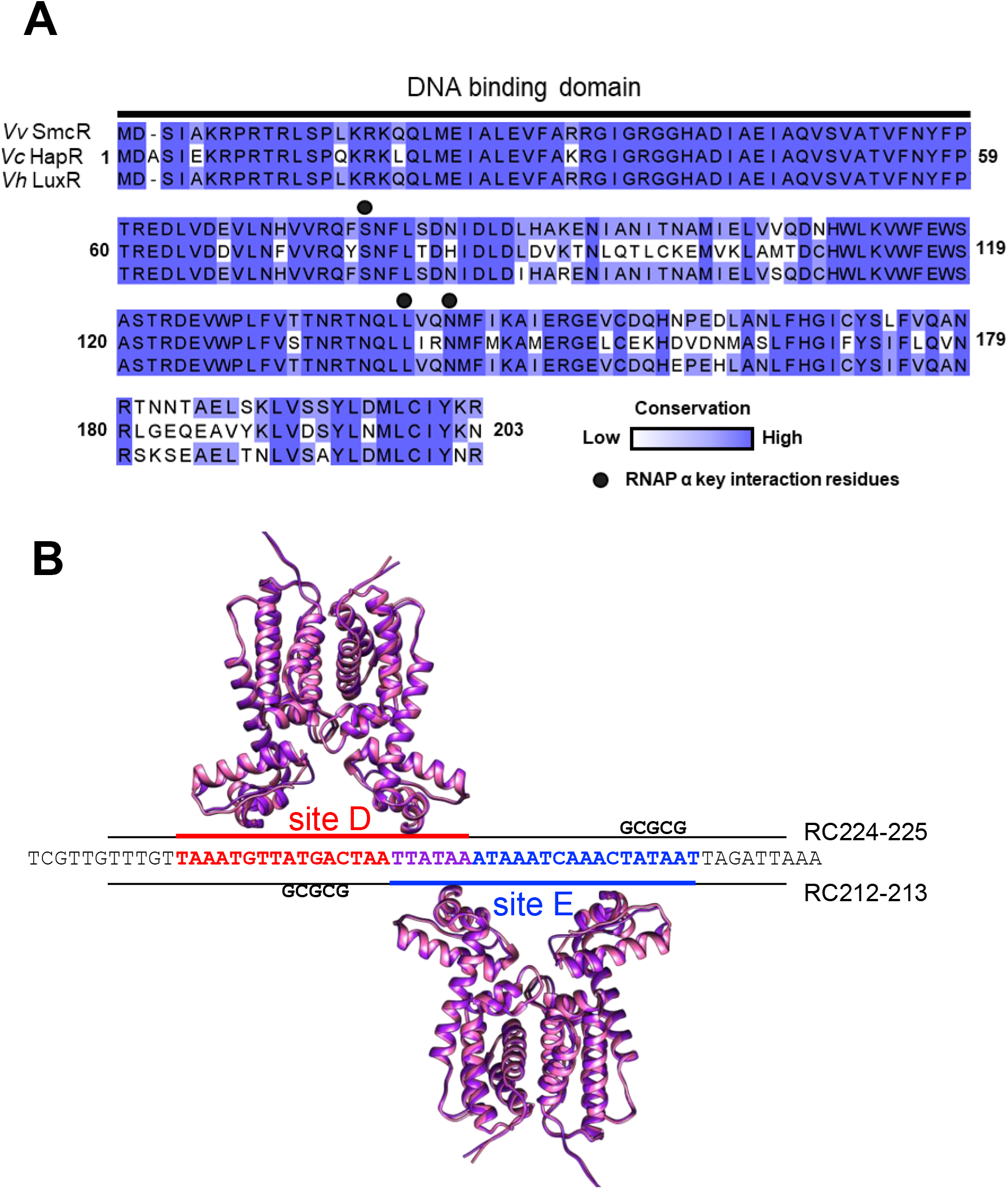
**(A)** Alignment of LuxR homologues from *V. vulnificus* (SmcR, CP012881), *V. harveyi* (LuxR, NC_009783) and *V. cholerae* (HapR, Q4F6W7) for residues 1 through 203, excluding the last three residues of SmcR and LuxR because HapR ends at 203. The DNA binding domain is indicated by a black line, and the residues (Ser76, Leu139, Asn142) previously shown to interact with RNA polymerase are indicated by a black circle above the residue. **(B)** Diagram of hypothesized SmcR binding orientation on two of the LuxR *V. harveyi* binding sites (D, red; E, blue; overlap, purple) that overlap from the *luxC* promoter. The changes made to oligonucleotides listed (RC212/213 and RC224/225) are shown in.

**Figure S2.**
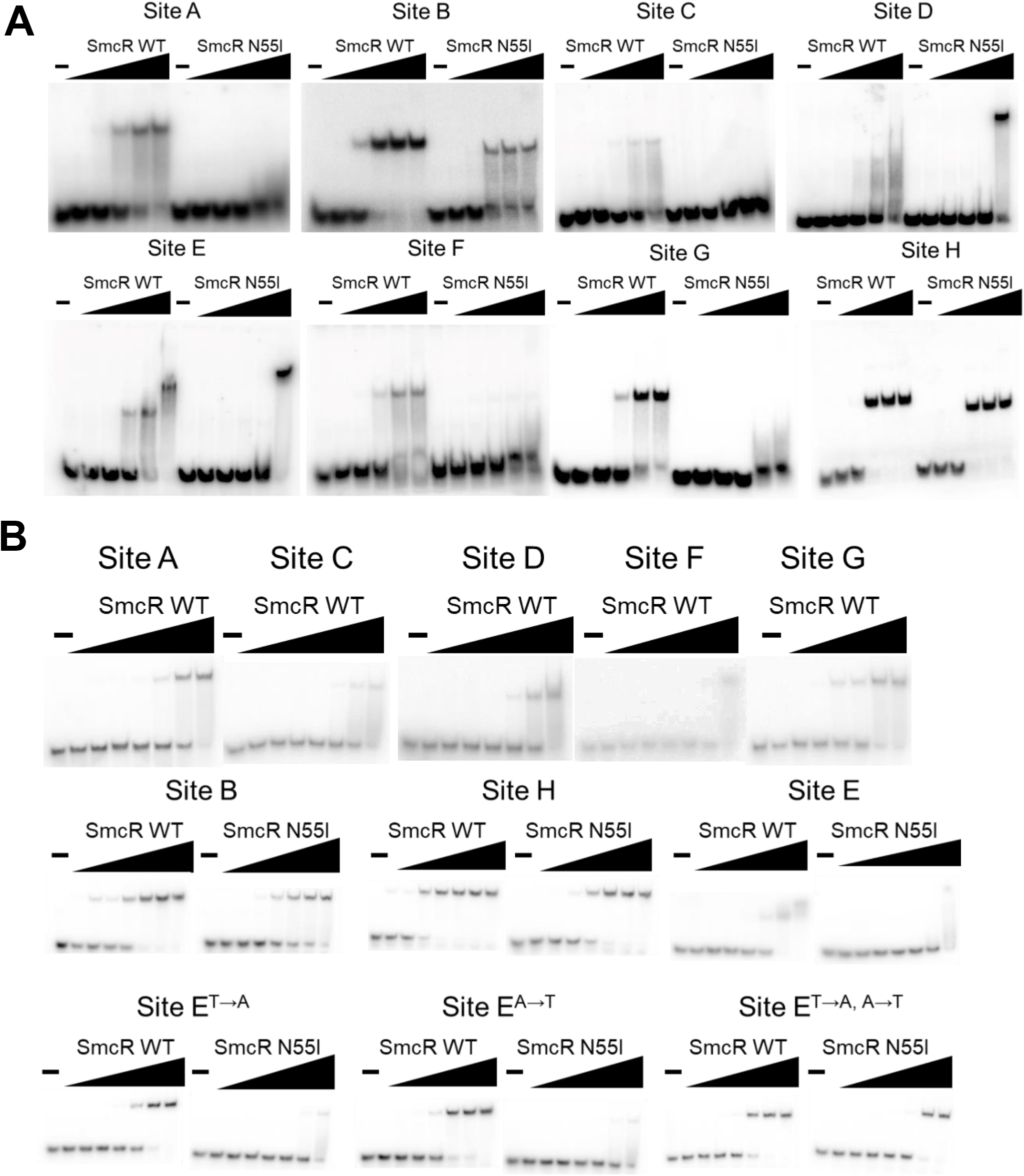
*(A)* EMSA reactions consisting of 0.5 nM radiolabeled DNA substrates from P_*luxC*_ (oligonucleotides listed in Table 1 and S3) and increasing concentrations of SmcR wild-type or SmcR N55I (0.1, 1, 10, 100, 1000 nM protein). Lanes labeled ‘—’ had no protein added. *(B)* EMSA reactions with 0.5 nM radiolabeled DNA from P_*luxC*_ sites and P_*luxC*_ substituted sites as indicated and increasing concentration SmcR wild-type or SmcR N55I as indicated (0.0005, 0.005, 0.05, 0.5, 5, 50, 500 nM protein). Lanes labeled ‘—’ had no protein added.

**Figure S3.**
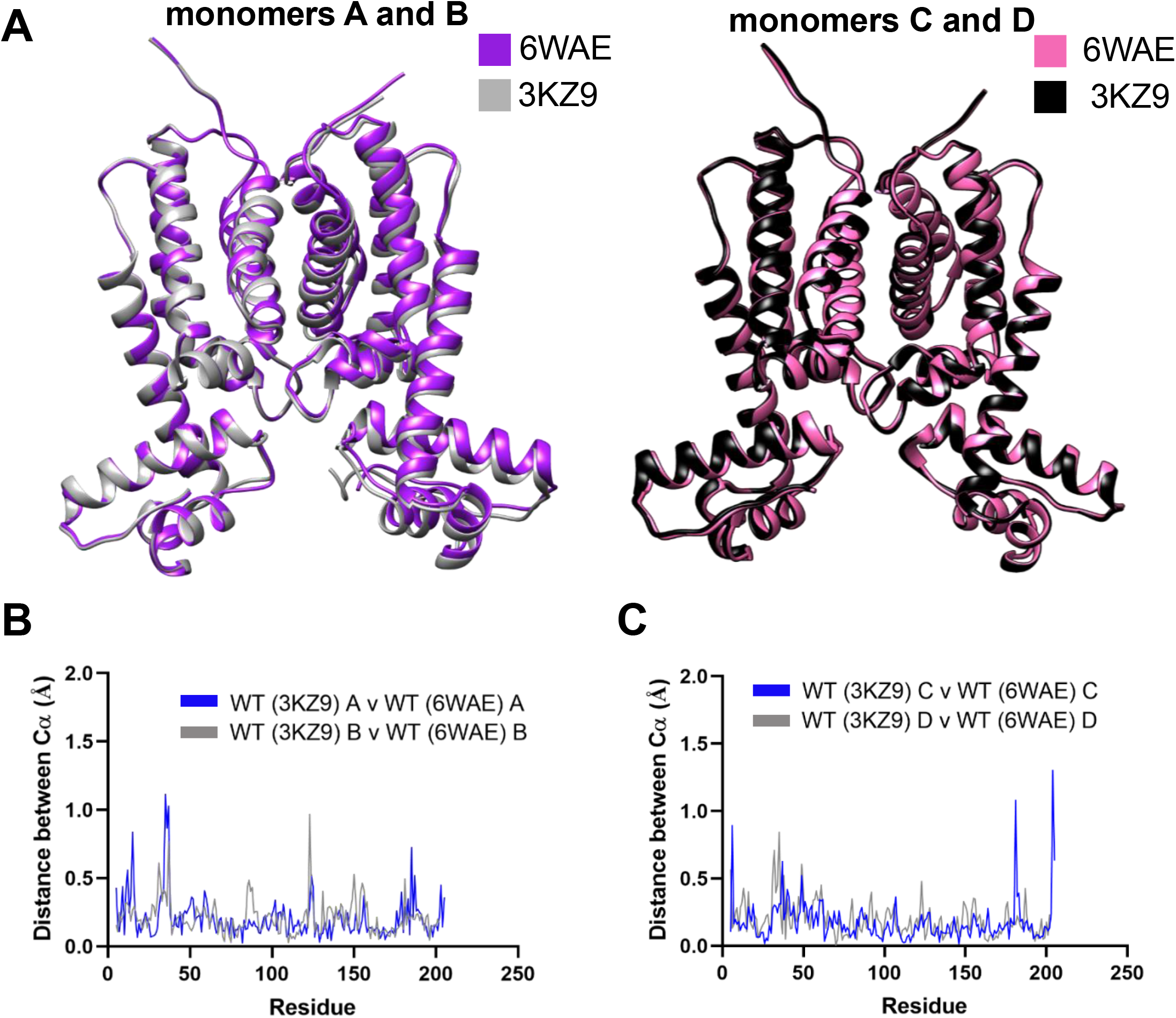
*(A, left)* Crystal structure of untagged SmcR wild-type (PDB: 3KZ9, gray) monomers A and B aligned with his-tagged SmcR wild-type (PDB: 6WAE, purple) monomers A and B. *(A, right)* Crystal structure of untagged SmcR wild-type (PDB: 3KZ9, black) monomers C and D aligned with his-tagged SmcR wild-type (PDB: 6WAE, pink) monomers C and D. *(B)* Graph showing the Chimera measured distances between alpha carbons (Cα) of untagged SmcR wild-type (PDB: 3KZ9) from his-tagged SmcR wild-type (PDB: 6WAE) for monomer A (blue) and monomer B (silver). *(C)* Graph showing Cα distances of SmcR wild-type (PDB: 3KZ9) from his-tagged SmcR wild-type (PDB: 6WAE) for monomer C (blue) and monomer D (silver).

**Figure S4.**
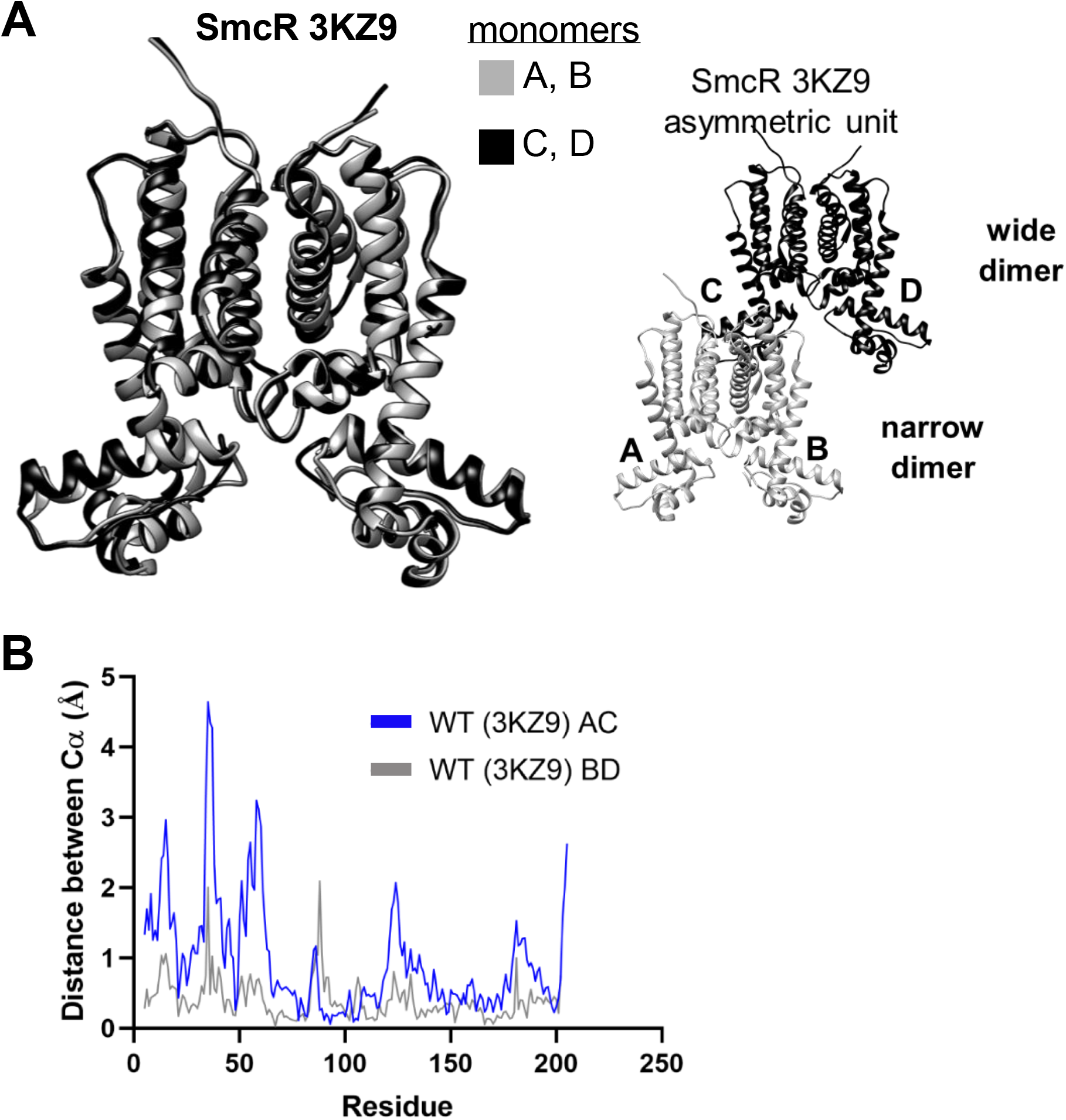
*(A, left)* Crystal structure of untagged SmcR wild-type (PDB: 3KZ9) monomers A and B (gray) aligned with monomers C and D (black). *(A, right)* The asymmetric unit containing two molecules of the wild-type untagged SmcR dimer (3KZ9; gray and black). *(B)* Graph showing the Chimera measured distances between alpha carbons (Cα) of his-tagged SmcR wild-type (PDB: 3KZ9) from monomer A to monomer C (gray) and from monomer B to monomer D (blue).

**Figure S5.**
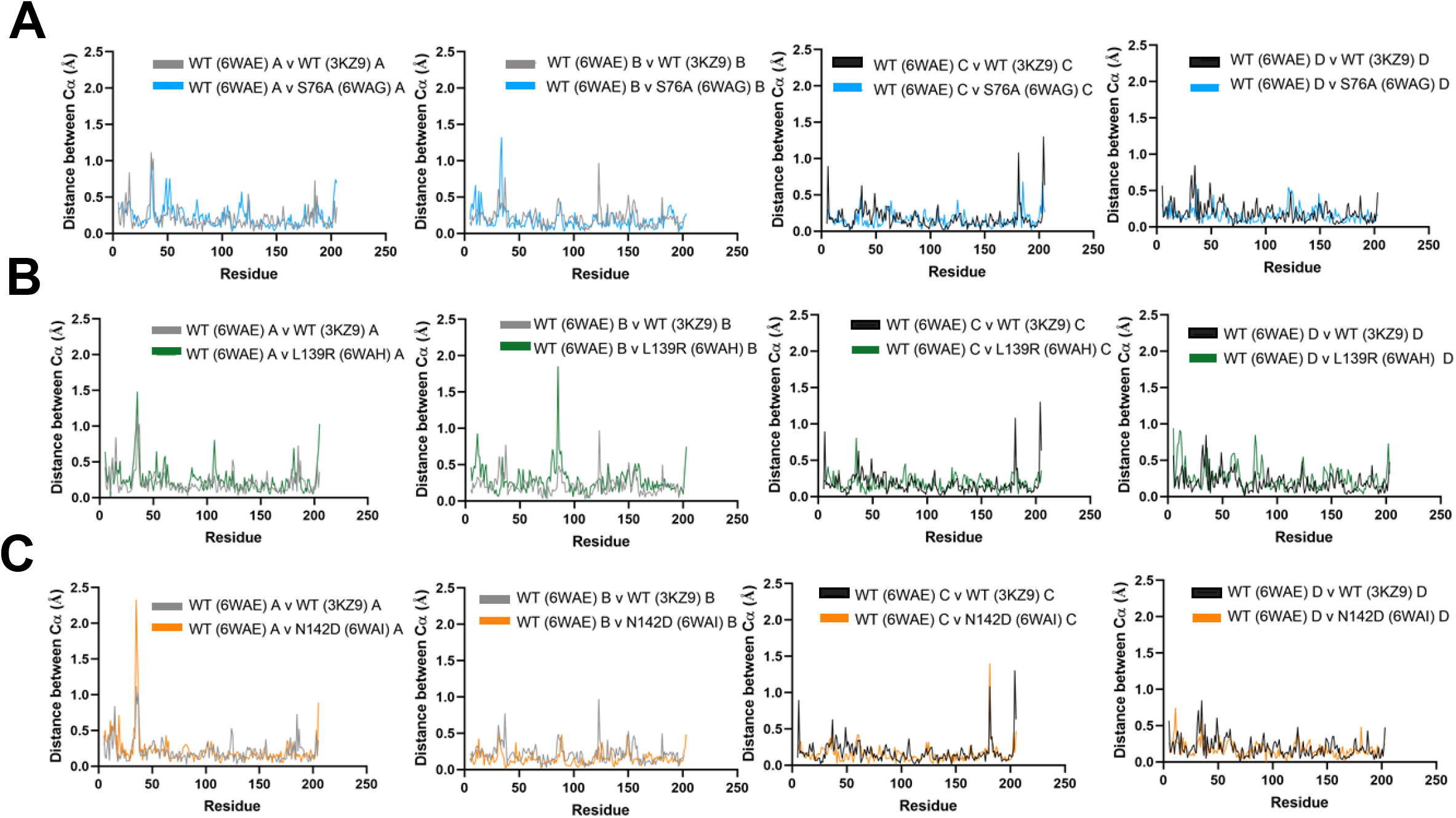
*(A)* Graphs comparing the distances between Cα of wild-type SmcR 6WAE monomers A-D vs SmcR S76A monomers A-D, respectively. *(B)* Graphs comparing the distance between Cα of wild-type SmcR 6WAE monomers A-D vs SmcR L139R monomers A-D, respectively. *(C)* Graphs comparing the distance between Cα of wild-type SmcR monomers A-D vs SmcR N142D monomers A-D, respectively.

**Figure S6.**
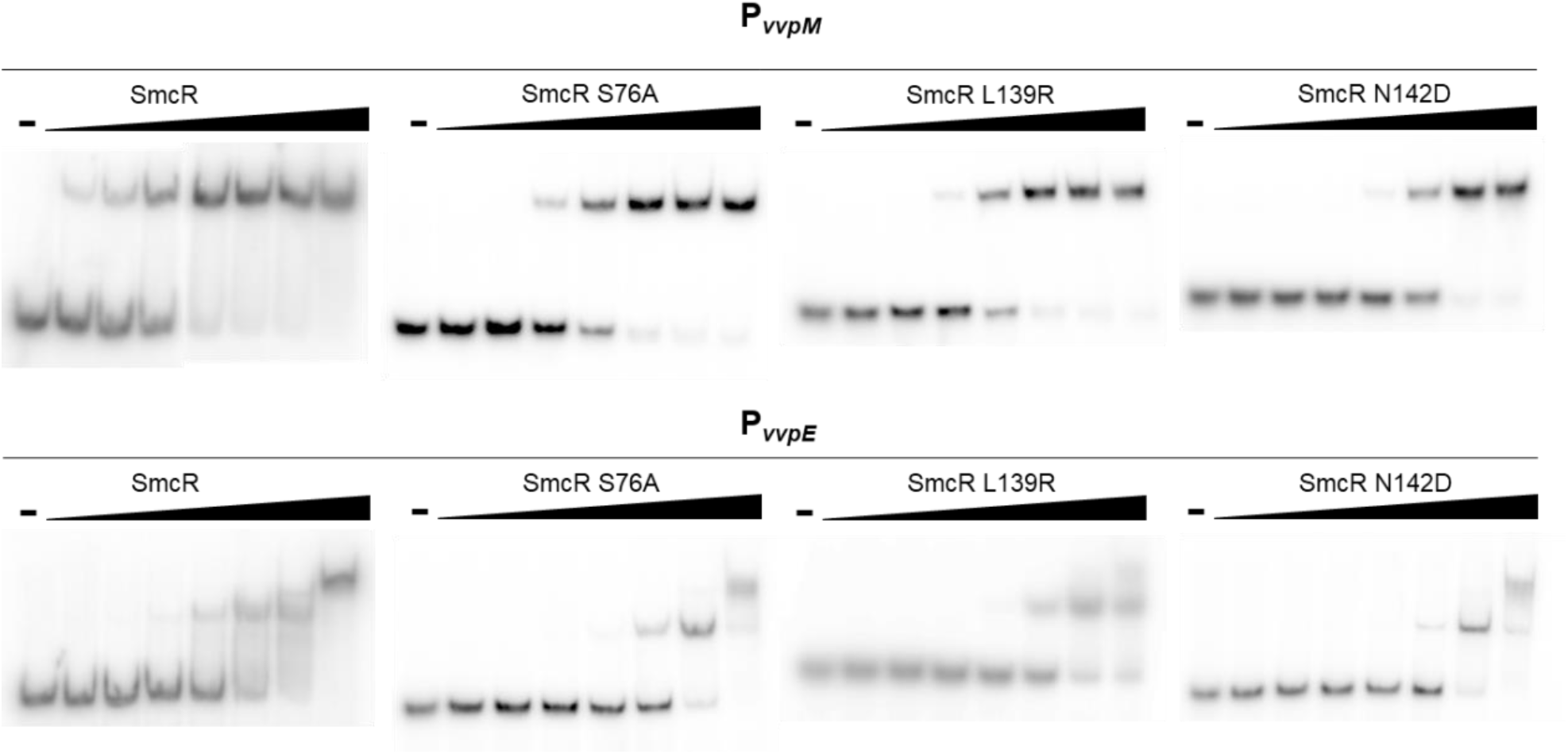
EMSA reactions consisting of 0.5 nM radiolabeled DNA substrates from P_*vvpM*_ or P_*vvpE*_ (oligonucleotides listed in Table S3) and increasing concentrations of SmcR wild-type or SmcR substitution mutants as listed (0.1, 1, 10, 100, 1000 nM protein). Lanes labeled ‘—’ had no protein added.

**Figure S7.**
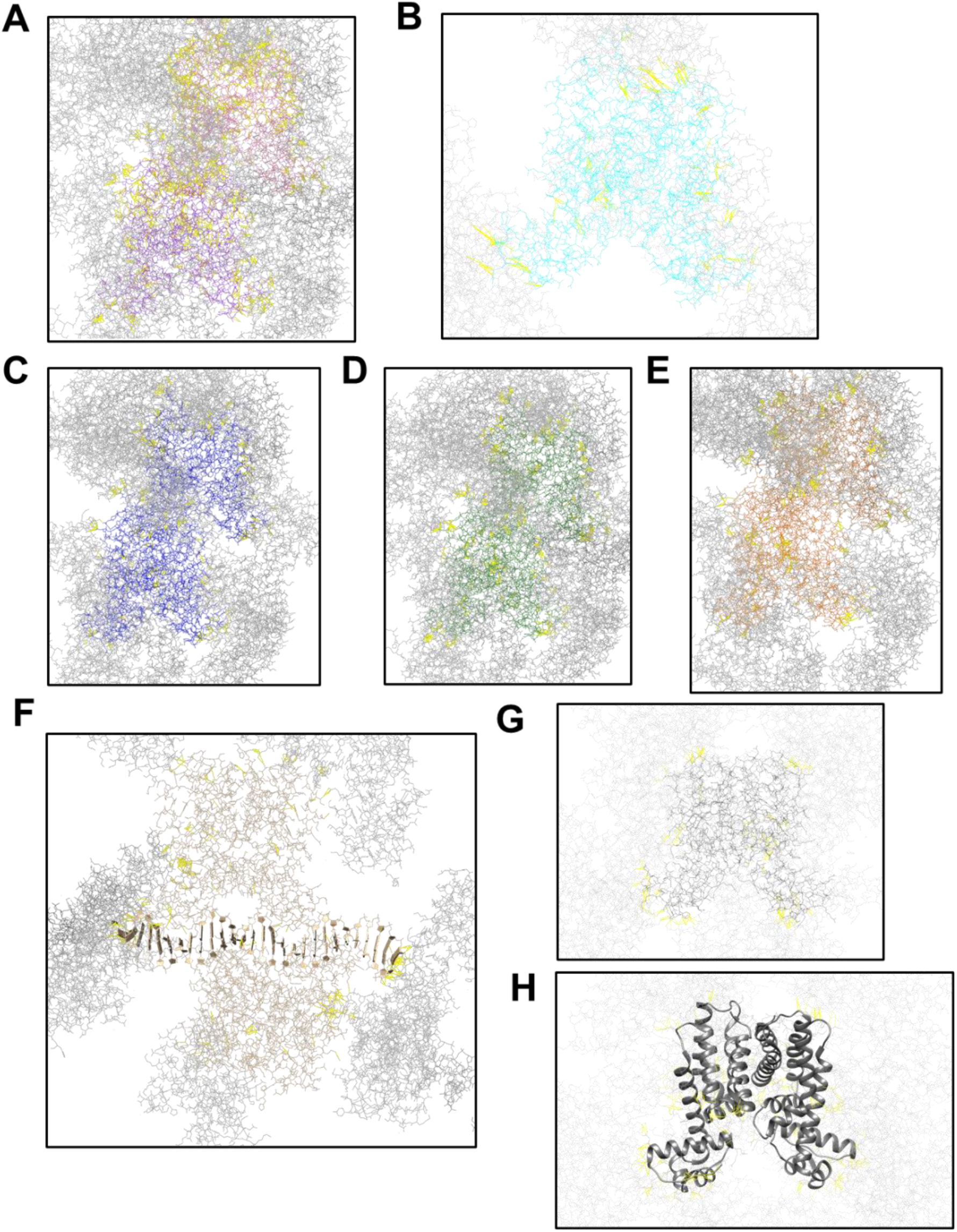
*(A)* Crystal contacts for SmcR WT (PDB: 6WAE) *(B)* Crystal contacts for SmcR N55I (PDB: 6WAF) *(C)* Crystal Contacts for SmcR S76A (PDB: 6WAG) *(D)* Crystal contacts for SmcR L139R (PDB: 6WAH) *(E)* Crystal contacts for SmcR N142D (PDB: 6WAI) *(F)* Crystal contacts for HapR (PDB: 2PBX) *(G)* Crystal contacts for QacR-DNA (PDB: 1JT0) *(H)* Crystal contacts for HapR_V2_ (PDB: 6D7R) with ribbon model for increased contrast.

**Table S1.**
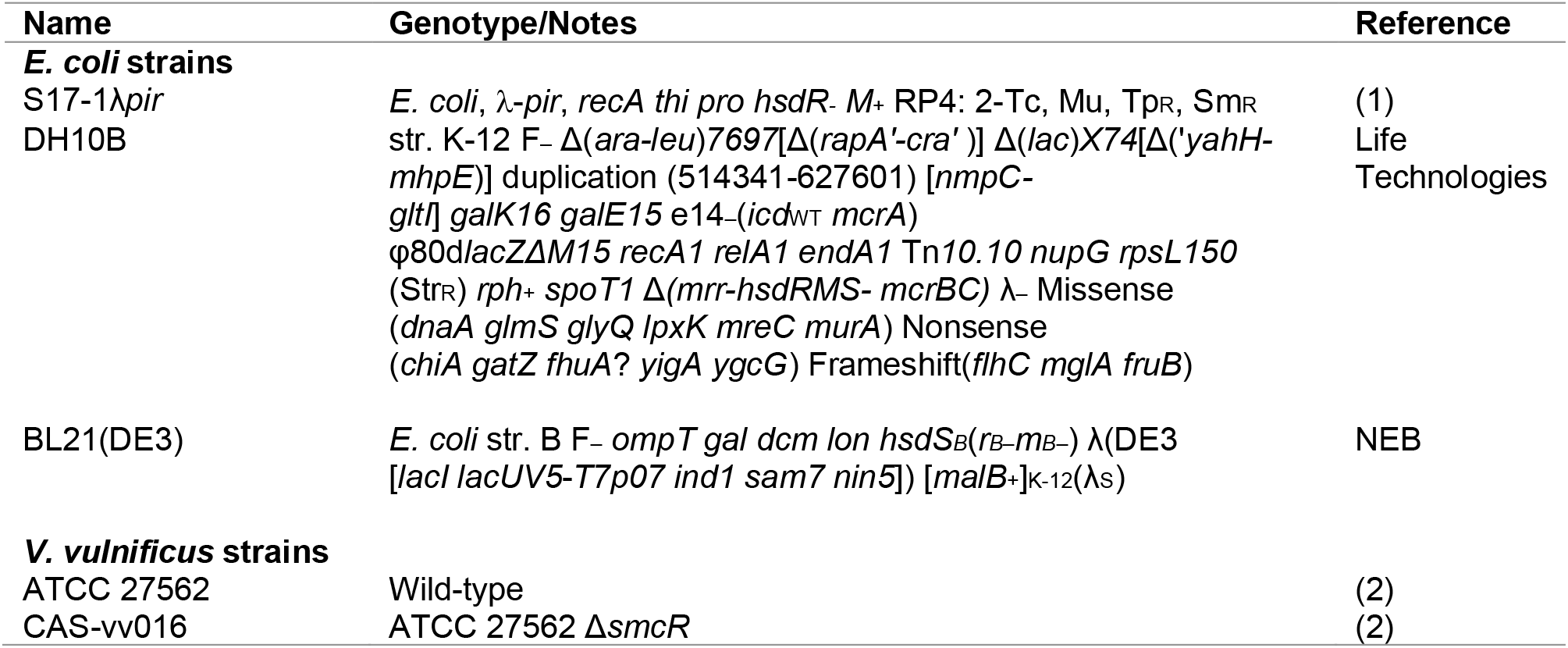
Strains used in this study.

**Table S2.**
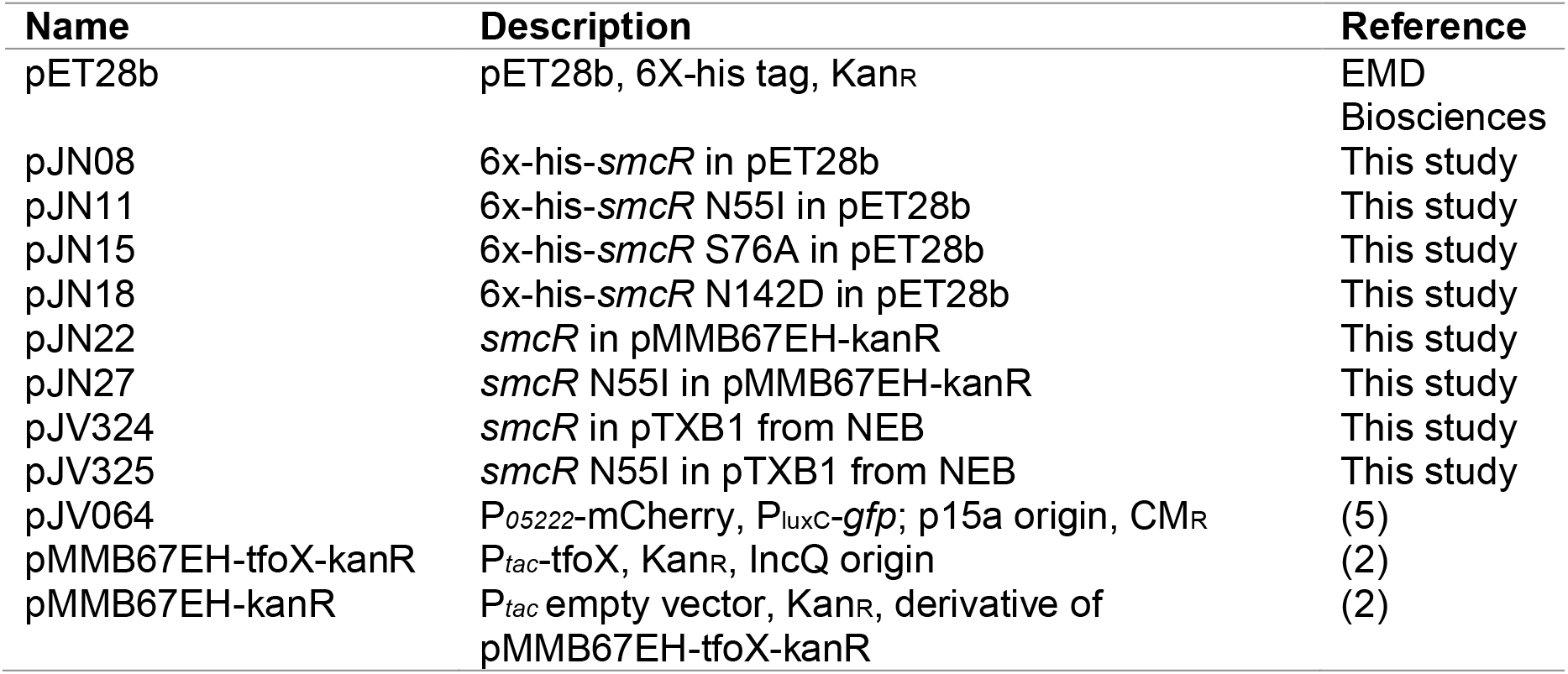
Plasmids used in this study.

| **Name** | **Description** | **Reference** |
| --- | --- | --- |
| pET28b | pET28b, 6X-his tag, Kan<sub>R</sub> | EMD<br>Biosciences |
| pJN08 | 6x-his-*smcR* in pET28b | This study |
| pJN11 | 6x-his-*smcR* N55I in pET28b | This study |
| pJN15 | 6x-his-*smcR* S76A in pET28b | This study |
| pJN18 | 6x-his-*smcR* N142D in pET28b | This study |
| pJN22 | *smcR* in pMMB67EH-kanR | This study |
| pJN27 | *smcR* N55I in pMMB67EH-kanR | This study |
| pJV324 | *smcR* in pTXB1 from NEB | This study |
| pJV325 | *smcR* N55I in pTXB1 from NEB | This study |
| pJV064 | P<sub>05222</sub>-mCherry, P<sub>luxC</sub>-*gfp*; p15a origin, CM<sub>R</sub> | (5) |
| pMMB67EH-tfoX-kanR | P<sub>tac</sub>-tfoX, Kan<sub>R</sub>, IncQ origin | (2) |
| pMMB67EH-kanR | P<sub>tac</sub> empty vector, Kan<sub>R</sub>, derivative of pMMB67EH-tfoX-kanR | (2) |

**Table S3.**
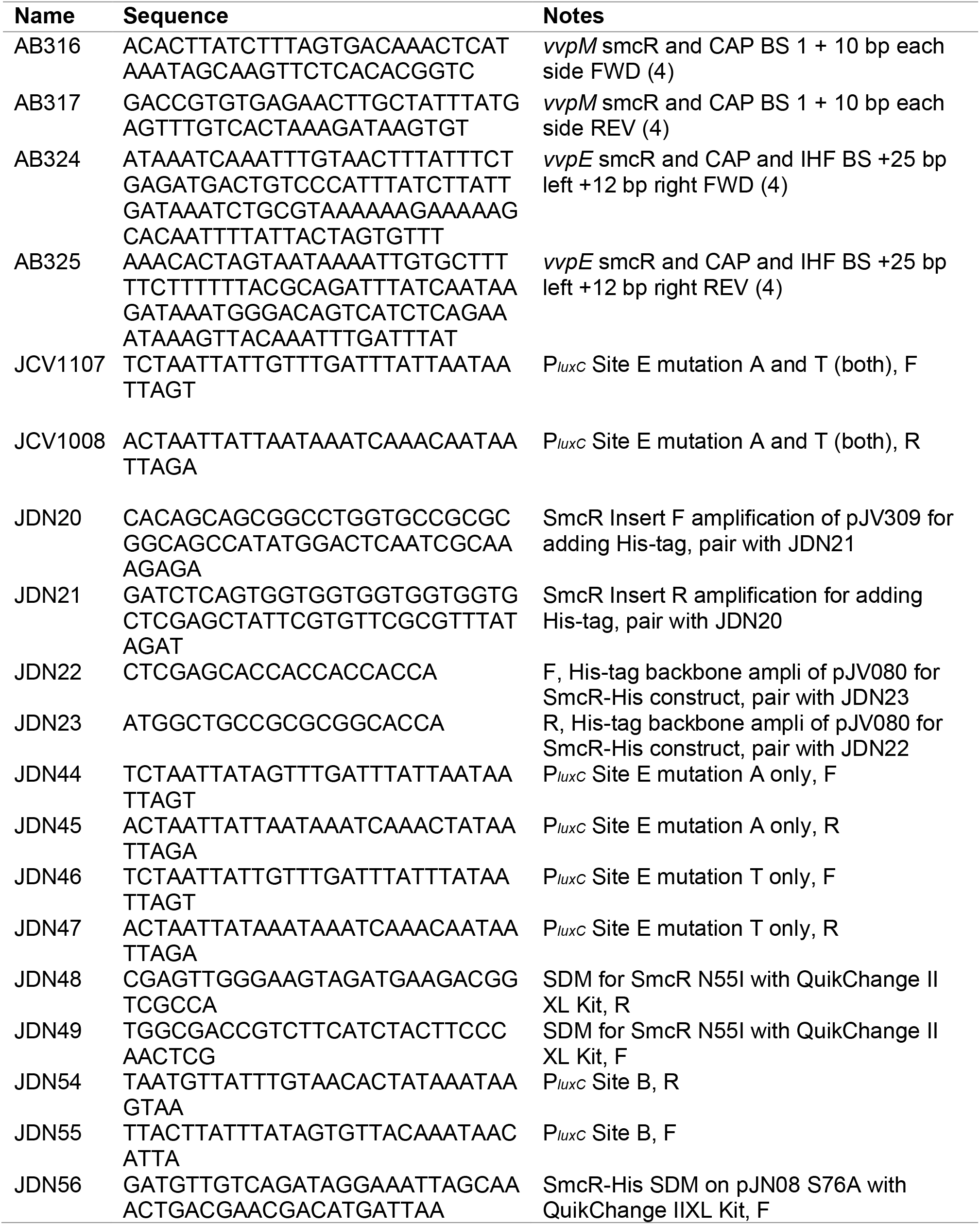
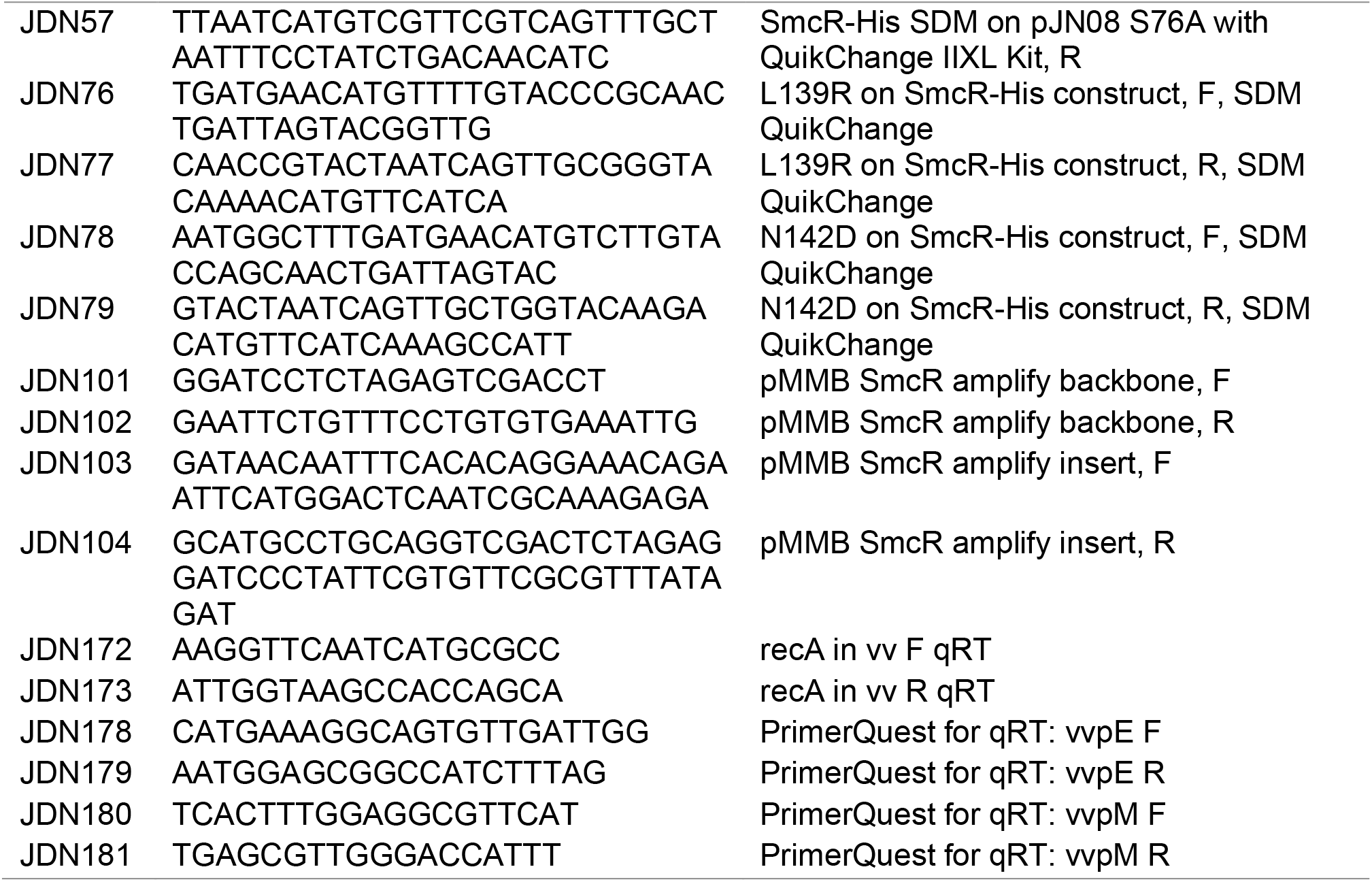
Oligonucleotides from this study.

**Table S4.** Crystallography data-collection and refinement statistics.

|  | **SmcR WT** | **SmcR N142D** | **SmcR S76A** | **SmcR L139R** | **SmcR N55I** |
| --- | --- | --- | --- | --- | --- |
| **Data collection** |  |  |  |  |  |
| Wavelength (Å) | 1.00003 | 0.97625 | 0.97625 | 1.00003 | 1.00003 |
| Space group | P212121 | P212121 | P212121 | P212121 | C2 |
| **Cell dimensions** |  |  |  |  |  |
| a, b, c (Å) | 78.16 99.32<br>129.53 | 78.02 99.56<br>129.47 | 78.04 99.04<br>129.68 | 77.97 99.00<br>129.11 | 167.57 57.92<br>78.11 |
| $\alpha, \beta, \gamma$ (°) | 90.00 90.00<br>90.00 | 90.00 90.00<br>90.00 | 90.00 90.00<br>90.00 | 90.00 90.00<br>90.00 | 90.00 109.82<br>90.00 |
| Resolution (Å) | 78.82 – 2.13<br>(2.21 – 2.13) | 78.92 – 2.58<br>(2.63 – 2.58) | 64.84 – 2.57<br>(2.62 – 2.57) | 46.22 – 2.55<br>(2.66 – 2.55) | 46.38 – 3.38<br>(3.65 – 3.38) |
| R<sub>sym</sub> | 0.137 (1.539) | 0.166 (1.623) | 0.158 (1.553) | 0.315 (1.265) | 0.141 (1.223) |
| R<sub>meas</sub> | 0.147 (1.688) | 0.179 (1.759) | 0.172 (1.694) | 0.373 (1.512) | 0.168 (1.476) |
| R<sub>pim</sub> | 0.055 (0.684) | 0.067 (0.673) | 0.067 (0.672) | 0.155 (0.588) | 0.090 (0.815) |
| Total reflections | 407960 (17017) | 229134 (10953) | 210347 (10120) | 219370 (25310) | 34296 (6667) |
| No. unique reflections | 57324 (2841) | 32305 (1592) | 32493 (1598) | 33302 (4024) | 9974 (2051) |
| CC1/2 | 0.999 (0.618) | 0.998 (0.626) | 0.998 (0.673) | 0.980 (0.786) | 0.996 (0.605) |
| I/$\sigma$(I) | 12.2 (1.1) | 10.1 (1.3) | 10.2 (1.3) | 7.7 (1.7) | 6.4 (1.1) |
| Completeness (%) | 100.0 (100.0) | 99.9 (100.0) | 99.9 (100.0) | 99.9 (99.9) | 98.9 (99.4) |
| Multiplicity | 7.1 (6.0) | 7.1 (6.9) | 6.5 (6.3) | 6.6 (6.3) | 3.4 (3.3) |
| Wilson B-factor | 35.26 | 42.34 | 43.54 | 32.24 | 99.22 |
| **Refinement** |  |  |  |  |  |
| Resolution (Å) | 66.95 – 2.13<br>(2.16 – 2.13) | 64.73 – 2.58<br>(2.66 – 2.58) | 55.43 – 1.95<br>(2.65 – 2.57) | 46.23 – 2.55<br>(2.63 – 2.55) | 41.00 – 3.38<br>(3.50 – 3.38) |
| No. unique reflections | 57219 (5628) | 32180 (3134) | 32242 (3166) | 33032 (3177) | 9317 (434) |
| R<sub>work</sub> | 0.2106 | 0.2313 | 0.2446 | 0.2280 | 0.2686 |
| R<sub>free</sub> | 0.2553 | 0.2860 | 0.2986 | 0.3031 | 0.2958 |
| CC(work) | 0.964 (0.819) | 0.956 (0.808) | 0.950 (0.803) | 0.950 (0.856) | 0.966 (0.535) |
| CC(free) | 0.960 (0.758) | 0.912 (0.719) | 0.902 (0.715) | 0.910 (0.736) | 0.922 (0.377) |
| **R.m.s.d values** |  |  |  |  |  |
| Bond lengths (Å) | 0.007 | 0.003 | 0.004 | 0.008 | 0.004 |
| Bond angles (°) | 0.703 | 0.662 | 0.633 | 0.856 | 0.700 |
| **No. atoms** |  |  |  |  |  |
| Protein | 6556 | 6521 | 6506 | 6550 | 3198 |
| Ligand/ions | 33 | 13 | 13 | 13 | 10 |
| ***B-factors*** (Å<sup>2</sup>) |  |  |  |  |  |
| Protein | 51.63 | 56.38 | 60.12 | 46.79 | 123.29 |
| Ligand/ions | 107.65 | 63.88 | 72.82 | 39.09 | 139.55 |
| ***Ramachandran plot*** |  |  |  |  |  |
| Favored (%) | 98.4 | 98.1 | 98.2 | 96.0 | 95.7 |
| Allowed (%) | 1.6 | 1.9 | 1.7 | 4.0 | 4.3 |
| Rotamer outliers (%) | 0.0 | 0.0 | 0.0 | 0.0 | 0.0 |
| *PDB code* | 6WAE | 6WAI | 6WAG | 6WAH | 6WAF |

\*Highest-resolution shell values are shown in parentheses.

## Notes

### Competing Interest Statement

The authors have declared no competing interest.

## References

1. M. B. Miller, K. Skorupski, D. H. Lenz, R. K. Taylor, B. L. Bassler, Parallel Quorum Sensing Systems Converge to Regulate Virulence in Vibrio cholerae. Cell 110, 303–314 (2002).

2. C. K. Ellison, et al., Retraction of DNA-bound type IV competence pili initiates DNA uptake during natural transformation in Vibrio cholerae. Nat. Microbiol. 3, 773–780 (2018).

3. R. T. Pena, et al., Relationship Between Quorum Sensing and Secretion Systems. Front. Microbiol. 10, 1100 (2019).

4. J. Henke, B. Bassler, Quorum sensing regulates type III secretion in Vibrio harveyi and Vibrio parahaemolyticus. J. Bacteriol. 186, 3794–3805 (2004).

5. D. L. McRose, et al., Quorum sensing and iron regulate a two-for-one siderophore gene cluster in Vibrio harveyi. PNAS 115 (2018).

6. R. Guillemette, B. Ushijima, M. Jalan, C. C. Häse, F. Azam, Insight into the resilience and susceptibility of marine bacteria to T6SS attack by Vibrio cholerae and Vibrio coralliilyticus. PLoS One 15, e0227864 (2020).

7. R. N. Paranjpye, M. S. Strom, Colonization of shellfish by pathogenic Vibrios in Proceedings of MTS/IEEE OCEANS, 2005, (2005).

8. S. A. Soto-Rodriguez, B. Gomez-Gil, R. Lozano-Olvera, M. Betancourt-Lozano, M. S. Morales-Covarrubias, Field and experimental evidence of Vibrio parahaemolyticus as the causative agent of acute hepatopancreatic necrosis disease of cultured shrimp (Litopenaeus vannamei) in northwestern Mexico. Appl. Environ. Microbiol. 81, 1689–1699 (2015).

9. L. Jayasree, P. Janakiram, R. Madhavi, Characterization of Vibrio spp. Associated with Diseased Shrimp from Culture Ponds of Andhra Pradesh (India). J. World Aquac. Soc. 37, 523–532 (2006).

10. H. M. Fadel, M. M. M. El-Lamie, Vibriosis and Aeromonas infection in shrimp: Isolation, sequencing, and control. Int. J. One Heal. 5, 38–48 (2019).

11. C. M. Waters, B. L. Bassler, Quorum Sensing : Communication in Bacteria. Annu. Rev. Cell Dev. Biol. 21, 319–346 (2005).

12. S. W. Teng, et al., Measurement of the copy number of the master quorum-sensing regulator of a bacterial cell. Biophys. J. 98, 2024–2031 (2010).

13. Z. Liu, A. Hsiao, A. Joelsson, J. Zhu, The Transcriptional Regulator VqmA Increases Expression of the Quorum-Sensing Activator HapR in Vibrio cholerae. J. Bacteriol. 188, 2446–2453 (2006).

14. S. M. Kim, et al., LuxR Homologue SmcR Is Essential for Vibrio vulnificus Pathogenesis and Biofilm Detachment, and Its Expression is Induced by Host Cells. Infect. Immun. 81, 3721–3730 (2013).

15. B. S. Kim, et al., QStatin, a Selective Inhibitor of Quorum Sensing inVibrioSpecies. MBio 9, e02262–17 (2018).

16. D. F. Browning, S. J. W. Busby, The Regulation of Bacterial Transcription Initiation. Nat. Rev. 2(2004).

17. S. J. Philips, et al., Allosteric transcriptional regulation via changes in the overall topology of the core promoter. Science (80-.). 349 (2018).

18. B. L. Jutras, A. Verma, B. Stevenson, Identification of Novel DNA-Binding proteins Using DNA-Affinity Chromatography/Pull Down. Curr. Protoc. Microbiol., 1–16 (2012).

19. J. L. Ramos, et al., The TetR family of transcriptional repressors. Microbiol. Mol. Biol. Rev. 69, 326–56 (2005).

20. D. H. Lee, et al., A Consensus Sequence for Binding of SmcR, a Vibrio vulnificus LuxR Homologue, and Genome-wide Identification of the SmcR Regulon * □ S. J. Biol. Chem. (2008) https:/doi.org/10.1074/jbc.M801480200 (March 21, 2018).

21. W. Lin, G. Kovacikova, K. Skorupski, Requirements for Vibrio cholerae HapR binding and transcriptional repression at the hapR promoter are distinct from those at the aphA promoter. J. Bacteriol. 187, 3013–9 (2005).

22. A. M. Tsou, T. Cai, Z. Liu, J. Zhu, R. V Kulkarni, Regulatory targets of quorum sensing in Vibrio cholerae: evidence for two distinct HapR-binding motifs. Nucleic Acids Res. 37, 2747–56 (2009).

23. J. C. van Kessel, L. E. Ulrich, I. B. Zhulin, B. L. Bassler, Analysis of activator and repressor functions reveals the requirements for transcriptional control by LuxR, the master regulator of quorum sensing in Vibrio harveyi. MBio 4, 1–10 (2013).

24. A. J. Pompeani, et al., The Vibrio harveyi master quorum-sensing regulator, LuxR, a TetR-type protein is both an activator and a repressor: DNA recognition and binding specificity at target promoters. Mol. Microbiol. 70, 76–88 (2008).

25. R. S. De Silva, et al., Crystal structure of the Vibrio cholerae quorum-sensing regulatory protein HapR. J. Bacteriol. 189, 5683–5691 (2007).

26. M. K. Jones, J. D. Oliver, Vibrio vulnificus: Disease and pathogenesis. Infect. Immun. 77, 1723–1733 (2009).

27. C.-M. Kim, et al., *Vibrio vulnificus* metalloprotease VvpE is essentially required for swarming. FEMS Microbiol. Lett. 269, 170–179 (2007).

28. A. S. Ball, J. C. van Kessel, The master quorum sensing regulators LuxR/HapR directly interact with the alpha subunit of RNA polymerase to drive transcription activation in *Vibrio harveyi* and *Vibrio cholerae*. Mol. Microbiol., 0–2 (2019).

29. H. S. Jeong, S. M. Kim, M. S. Lim, K. S. Kim, S. H. Choi, Direct interaction between quorum-sensing regulator SmcR and RNA polymerase is mediated by integration host factor to activate vvpE encoding elastase in Vibrio vulnificus. J. Biol. Chem. (2010) https:/doi.org/10.1074/jbc.M109.089987.

30. R. R. Chaparian, S. G. Olney, C. M. Hustmyer, D. A. Rowe-Magnus, J. C. van Kessel, Integration host factor and LuxR synergistically bind DNA to coactivate quorum-sensing genes in Vibrio harveyi. Mol. Microbiol. 101, 823–840 (2016).

31. M.-A. Lee, et al., VvpM, an extracellular metalloprotease of Vibrio vulnificus, induces apoptotic death of human cells. J. Microbiol. 52, 1036–1043 (2014).

32. C. P. Shao, L. I. Hor, Regulation of metalloprotease gene expression in Vibrio vulnificus by a Vibrio harveyi LuxR homologue. J. Bacteriol. 183, 1369–75 (2001).

33. Y. Kim, et al., Crystal structure of SmcR, a quorum-sensing master regulator of Vibrio vulnificus, provides insight into its regulation of transcription. J. Biol. Chem. 285, 14020–14030 (2010).

34. M. Dongre, et al., Evidence on How a Conserved Glycine in the Hinge Region of HapR Regulates Its DNA Binding Ability LESSONS FROM A NATURAL VARIANT * □ S Downloaded from. J. Biol. Chem. 286, 15043 (2011).

35. T. A. Azam, A. Ishihama, Twelve species of the nucleoid-associated protein from Escherichia coli. Sequence recognition specificity and DNA binding affinity. J. Biol. Chem. 274, 33105–33113 (1999).

36. R. R. Chaparian, M. L. N. Tran, L. C. Miller Conrad, D. B. Rusch, J. C. Van Kessel, Global H-NS counter-silencing by LuxR activates quorum sensing gene expression. Nucleic Acids Res. 48, 171–183 (2020).

37. G. Brackman, et al., Structure-activity relationship of cinnamaldehyde analogs as inhibitors of AI-2 based quorum sensing and their effect on virulence of Vibrio spp. PLoS One 6(2011).

38. M. J. Kratochvil, T. Yang, H. E. Blackwell, D. M. Lynn, Nonwoven Polymer Nanofiber Coatings That Inhibit Quorum Sensing in Staphylococcus aureus: Toward New Nonbactericidal Approaches to Infection Control. ACS Infect. Dis. (2017) https:/doi.org/10.1021/acsinfecdis.6b00173 (April 25, 2017).

39. G. Brackman, et al., Cinnamaldehyde and cinnamaldehyde derivatives reduce virulence in Vibrio spp. by decreasing the DNA-binding activity of the quorum sensing response regulator LuxR. BMC Microbiol. 8, 149 (2008).

40. T. Defoirdt, et al., A quorum sensing-disrupting brominated thiophenone with a promising therapeutic potential to treat luminescent vibriosis. PLoS One 7, 1–7 (2012).

41. J. Cruite, P. Succo, S. Raychaudhuri, F. J. Kull, Crystal structure of an inactive variant of the quorum-sensing master regulator HapR from the protease-deficient non-O1, non-O139 Vibrio cholerae strain V2. Acta Crystallogr. Sect. F, Struct. Biol. Commun. 74, 331–336 (2018).

42. B. Lin, et al., Comparative genomic analyses identify the Vibrio harveyi genome sequenced strains BAA-1116 and HY01 as Vibrio campbellii. Environ. Microbiol. Rep. 2, 81–89 (2010).

43. S. T. Rutherford, J. C. Van Kessel, Y. Shao, B. L. Bassler, AphA and LuxR/HapR reciprocally control quorum sensing in vibrios. Genes Dev. (2011) https:/doi.org/10.1101/gad.2015011.

44. J. D. Newman, J. C. van Kessel, Purification of the Vibrio Quorum-Sensing Transcription Factors LuxR, HapR, and SmcR. Methods Mol. Biol. (2020) https:/doi.org/10.1007/7651_2020_306.

45. W. Kabsch, et al., XDS. Acta Crystallogr. Sect. D Biol. Crystallogr. 66, 125–132 (2010).

46. P. Emsley, B. Lohkamp, W. G. Scott, K. Cowtan, Features and development of Coot. Acta Crystallogr. Sect. D Biol. Crystallogr. 66, 486–501 (2010).

47. P. D. Adams, et al., PHENIX: A comprehensive Python-based system for macromolecular structure solution. Acta Crystallogr. Sect. D Biol. Crystallogr. 66, 213–221 (2010).

48. E. F. Pettersen, et al., UCSF Chimera - A visualization system for exploratory research and analysis. J. Comput. Chem. 25, 1605–1612 (2004).

49. L. Schrödinger, The PyMOL Molecular Graphics System, Version 2.0 (2015).

## Supporting References

1. V. de Lorenzo, K. N. Timmis, Analysis and construction of stable phenotypes in gramnegative bacteria with Tn5- and Tn10-derived minitransposons. Methods Enzymol. 235, 386–405 (1994).

2. C. A. Simpson, R. Podicheti, D. B. Rusch, A. B. Dalia, J. C. van Kessel, Diversity in natural transformation frequencies and regulation across vibrio species. MBio 10 (2019).

3. J. C. Van Kessel, S. T. Rutherford, Y. Shao, A. F. Utria, B. L. Bassler, Individual and combined roles of the master regulators apha and luxr in control of the Vibrio harveyi quorum-sensing regulon. J. Bacteriol. 195, 436–443 (2013).

4. A. S. Ball, J. C. van Kessel, The master quorum sensing regulators LuxR/HapR directly interact with the alpha subunit of RNA polymerase to drive transcription activation in *Vibrio harveyi* and *Vibrio cholerae*. Mol. Microbiol., 0–2 (2019).

5. J. C. van Kessel, L. E. Ulrich, I. B. Zhulin, B. L. Bassler, Analysis of activator and repressor functions reveals the requirements for transcriptional control by LuxR, the master regulator of quorum sensing in Vibrio harveyi. MBio 4, 1–10 (2013).

